# From primordial clocks to circadian oscillators

**DOI:** 10.1101/2022.11.28.518275

**Authors:** Warintra Pitsawong, Ricardo A. P. Pádua, Timothy Grant, Marc Hoemberger, Renee Otten, Niels Bradshaw, Nikolaus Grigorieff, Dorothee Kern

## Abstract

Circadian rhythms play an essential role in many biological processes and surprisingly only three prokaryotic proteins are required to constitute a true post-translational circadian oscillator. The evolutionary history of the three Kai proteins indicates that KaiC is the oldest member and central component of the clock, with subsequent additions of KaiB and KaiA to regulate its phosphorylation state for time synchronization. The canonical KaiABC system in cyanobacteria is well understood, but little is known about more ancient systems that possess just KaiBC, except for reports that they might exhibit a basic, hourglass-like timekeeping mechanism. Here, we investigate the primordial circadian clock in *Rhodobacter sphaeroides* (RS) that contains only KaiBC to elucidate its inner workings despite the missing KaiA. Using a combination X-ray crystallography and cryo-EM we find a novel dodecameric fold for KaiC_RS_ where two hexamers are held together by a coiled-coil bundle of 12 helices. This interaction is formed by the C-terminal extension of KaiC_RS_ and serves as an ancient regulatory moiety later superseded by KaiA. A coiled-coil register shift between daytime- and nighttime-conformations is connected to the phosphorylation sites through a long-range allosteric network that spans over 160 Å. Our kinetic data identify the difference in ATP-to-ADP ratio between day and night as the environmental cue that drives the clock and further unravels mechanistic details that shed light on the evolution of self-sustained oscillators.

## Main Text

Circadian clocks are self-sustained biological oscillators that are found ubiquitously in prokaryotic and eukaryotic organisms. In eukaryotes these systems are complex and very sophisticated, whereas in prokaryotes the core mechanism is regulated by a posttranslational oscillator that can be reconstituted *in vitro* with three proteins (*kaiA, kaiB*, and *kaiC*)^1^ and ATP. Seminal work on the KaiABC system resulted in a comprehensive understanding of its circadian clock: KaiC is the central component that auto-phosphorylates through binding of KaiA and auto-dephosphorylates upon association with KaiB^2-5^. The interplay between these three proteins has been shown *in vitro* to constitute a true circadian oscillator characterized by persistence, resetting, and temperature compensation. Consequently, the KaiABC system is considered an elegant and the simplest implementation of a circadian rhythm. The evolutionary history of *kai* genes established *kaiC* as the oldest member dating back ∼3.5 bya, with subsequent additions of *kaiB* and most recently *kaiA* to form the extant *kaiBC* and *kaiABC* clusters, respectively^6,7^. Interestingly, a few studies of more primitive organisms that lack *kaiA* hinted that the *kaiBC*-based systems might provide already a basic, hourglass-like timekeeping mechanism^8-10^. Contrary to the self-sustained oscillators found in cyanobacteria, such a timer requires an environmental cue to drive the clock and for the daily “flip” of the hourglass. The central role of circadian rhythms in many biological processes, controlled by the day and night cycle on earth, makes their evolution a fascinating topic.

Here, we investigate such a primitive circadian clock by biochemical and structural studies of the KaiBC system of the purple, non-sulfur photosynthetic proteobacterium *Rhodobacter sphaeroides* KD131 (RS; hereafter referred to as KaiB_RS_ and KaiC_RS_). Surprisingly, the organism shows sustained rhythms of gene expression *in vivo*, but whether *kaiBC* is responsible for this observation remains inconclusive in the absence of a *kaiC* knockout^11^. A more recent study of the closely-related *Rhodopseudomonas palustris* demonstrated causality between the proto-circadian rhythm of nitrogen fixation and expression of the *kaiC* gene using a KO strain^10^. We discover using *in vitro* experiments that KaiBC_RS_ is indeed a primordial circadian clock with a mechanism that is different from the widely studied circadian oscillator in *Synechococcus elongatus* PCC 7942 (hereafter referred to as KaiABC_SE_)^2-5^. We identify an environmental cue that regulates the phosphorylation state and consequently produces a 24-hour clock *in vivo* as the switch in ATP-to-ADP ratio between day and night. Our kinetic results combined with X-ray and cryo-EM structures of the relevant states unravel a long-range allosteric pathway that is crucial for function of the hourglass and sheds light on the evolution of self-sustained oscillators. Notably, we find a novel protein fold for KaiC_RS_ and uncover a register shift in the coiled-coil domain spanning ∼115 Å as the key regulator in this system, which shows intriguing structural similarities to dynein signaling^12^.

### C-terminal tail as a primitive regulatory moiety

To gain insight into the evolution of the *kaiBC* cluster, we constructed a phylogenetic tree of *kaiC* after the emergence of the *kaiB* gene (Fig. 1a, Extended Data Fig. 1a). The first obvious question is how KaiC_RS_ and other members in the clade can auto-phosphorylate despite having no KaiA, which is known to be crucial for this function in the canonical KaiABC system at its optimum temperature. We observe a large clade that exhibits a C-terminal tail about 50 amino acids longer compared to *kaiC* in other clades (Extended Data Fig. 1b). This C-terminal extension near the A-loop is predominantly found in the *kaiC2* subgroup, which was previously annotated as having two serine phosphorylation sites instead of the Thr/Ser pair found in *kaiC1* and *kaiC3* subgroups (Extended Data Fig. 1b)^13-15^. In *Synechococcus elongatus*, the binding of KaiA_SE_ to the A-loop of KaiC_SE_ tethers them in an exposed conformation^16^ that activates both auto-phosphorylation and nucleotide exchange^17^. Given the proximity of the extended C-terminal tail to the A-loop we conjectured that it could serve as the “primitive” regulatory moiety made redundant concomitantly with the appearance of KaiA.

**Fig. 1.**
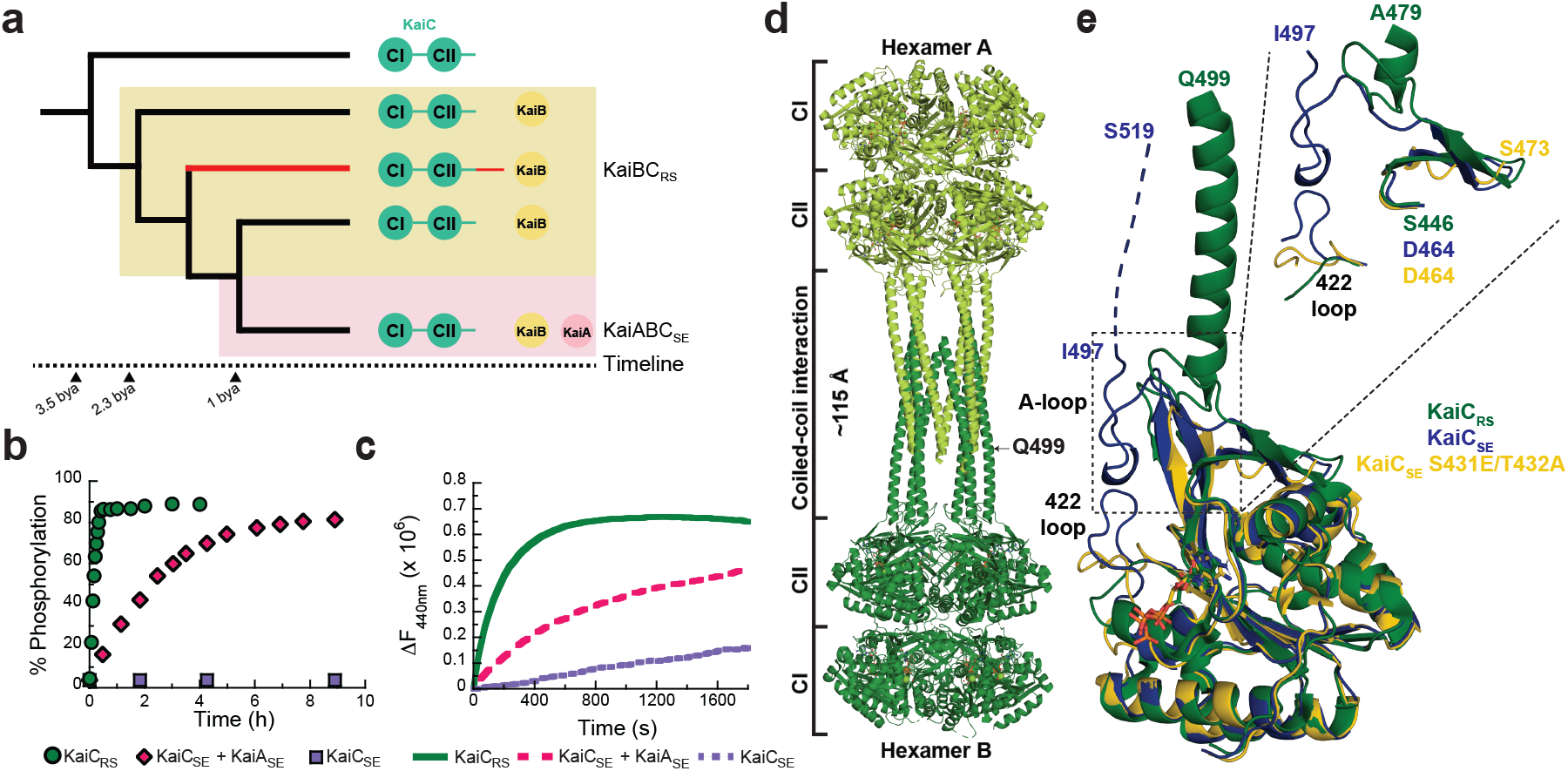
Extended C-terminal tail of KaiC_RS_ forms a coiled-coil interaction with an exposed A-loop for KaiA-independent phosphorylation of KaiC. **(a)** Schematic phylogenetic tree of *kaiC* showing the appearance of *kaiB* and *kaiA* during evolution. The *kaiC* clade with an approximately 50 amino acids C-terminal extension is labeled in red and a timeline was predicted as previously reported^6^ **(b)** Phosphorylation over time of KaiC_RS_ (green circles, 6.5 ± 1.0 h^-1^) and KaiC_SE_ in the presence (pink circles, 0.40 ± 0.02 h^-1^) or absence (purple circles) of KaiA_SE_ at 30 °C. The standard deviation in reported parameters are obtained from the fitting. **(c)** Nucleotide exchange between ATP and mant-ATP in KaiC_RS_ alone (green trace, 18.0 ± 1.5 h^-1^) compared to KaiC_SE_ in the presence (pink trace, 4.7 ± 0.3 h^-1^) and absence of KaiA_SE_ (purple trace, 0.08 ± 0.04 h^-1^) measured by fluorescence at 30 °C. Representative traces are shown and the fitted parameters (mean ± S.D) were obtained from three replicate measurements. **(d)** X-ray structure of dodecameric KaiC_RS_ (PDB 8dba) colored by hexamer A (light green) and hexamer B (dark green). The CI, CII, and coiled-coil domains are indicated in (a), and the A-loop is labeled in (e). **(e)** Superposition based on alignment of the CII domain of KaiC_RS_ (green, PDB 8dba, chain B), KaiC_SE_ (purple, PDB 1tf7, chain B)^36^, and KaiC_SE_ S431E/T432A (yellow, PDB 7s65, chain A)^19^ shows that KaiC_RS_ has an extended A-loop orientation that no longer forms the inhibitory interaction with the 422-loop (KaiC_SE_ numbering). The conformation of the 422-loop in KaiC_RS_ resembles the one seen in the cryo-EM structure of the phosphomimetic KaiC_SE_ S431E/T432A (yellow, PDB 7s65)^19^. No electron density is observed for the C-terminal part of wild-type KaiC_SE_ and S431E/T432A mutant due to flexibility, and the missing 22 residues for wild-type KaiC_SE_ (46 for S431E/T432A) are represented by a dashed line for wild-type KaiC_SE_ (not shown for the mutant).

To test our hypothesis, we first measured the auto-phosphorylation and nucleotide exchange rates in KaiC_RS_ that both depend heavily on the presence of KaiA in the KaiABC_SE_ system. We observe an auto-phosphorylation rate for KaiC_RS_ that is ∼16-fold higher than for KaiC_SE_ activated by KaiA_SE_ (6.5 ± 1.0 h^-1^ versus 0.40 ± 0.02 h^-1^, respectively; Fig. 1b and Extended Data Figs. 2a-e). Similarly, the nucleotide exchange is faster in KaiC_RS_ compared to KaiC_SE_ even in the presence of KaiA_SE_ (18.0 ± 1.5 h^-1^ compared to 4.7 ± 0.3 h^-1^, respectively; Fig. 1c and Extended Data Fig. 2f). Our data clearly show that KaiC_RS_ can perform both auto-phosphorylation and nucleotide exchange on its own and, in fact, does so faster than its more recently evolved counterparts.

### Coiled-coil interaction assembles KaiC_RS_ dodecamer

To assess mechanistically how KaiC in the *kaiA*-lacking systems accomplishes auto-phosphorylation we turned to structural biology. The crystal structure of KaiC_RS_, unlike KaiC from cyanobacteria, reveals a homododecamer consisting of two homohexameric domains joined by a twelve-helical coiled-coil domain that is formed by the extended C-terminal tail (PDB 8dba; Fig. 1d and Extended Data Table 1). A closer inspection of the CII domains in KaiC_RS_ and KaiC_SE/TE_ showed an obvious difference in A-loop orientations: an extended conformation in KaiC_RS_ versus a buried orientation in KaiC_SE/TE_ (Fig. 1e). The existence of such an extended conformation upon binding of KaiA was conjectured earlier based on the perceived hyper- and hypo-phosphorylation upon removing the A-loop or disrupting KaiA binding, respectively^18^. Importantly, a recent cryo-EM structure of the nighttime phosphomimetic KaiC_SE_ S431E/T432A in its compressed state was solved and directly showed a disordered A-loop that no longer interacts with the 422-loop^19^, similar to the extended A-loop conformation we observe in KaiC_RS_ (Fig. 1e). The loss of interaction between the A-loop and 422-loop (just 10 residues apart from the phosphorylation sites), results in closer proximity between the hydroxyl group of Ser^431^/Thr^432^ and the γ-phosphate of ATP, thereby, facilitating the phosphoryl-transfer step^20^. Furthermore, the sequence similarity between KaiC_RS_ and KaiC_SE_ is less than 30% for the A-loop and residues considered important for stabilization of this loop in its buried orientation (i.e., 422-loop and residues 438-444; Fig. 1e). Taken together, our structural and kinetic data support the idea that an exposed A-loop is key for KaiA-independent enhancement of nucleotide exchange and hence auto-phosphorylation in KaiC_RS_ and perhaps other KaiBC-based systems.

Is the purpose of the coiled-coil domain merely to “pull up” the A-loop or does it actively participate in nucleotide exchange and auto-phosphorylation of KaiC? To further understand its role, we generated a truncation at residue Glu^490^ based on the phylogenetic tree and crystallographic information (KaiC_RS_-Δcoil; Extended Data Fig. 1b) to disrupt the coiled-coil interaction between the two hexamers. Indeed, the crystal structure of KaiC_RS_-Δcoil (PDB 8db3; Fig. 2a,b and Extended Data Table 1) and its size-exclusion chromatogram and analytical ultracentrifugation profile (Extended Data Fig. 3a-c) show a hexameric structure with no coiled-coil interaction. Nucleotide exchange rates in the CII domain for KaiC_RS_-Δcoil and wild type are comparable (19.1 ± 0.8 h^-1^ and 18.0 ± 1.5 h^-1^, respectively; Extended Data Fig. 3d) as are the phosphorylation rates (5.5 ± 0.4 h^-1^ and 7.4 ± 0.3 h^-1^ for KaiC_RS_-Δcoil and wild type, respectively; Extended Data Fig. 3e,f). These results indicate that the extended A-loop and not the coiled-coil interaction plays a pivotal role in nucleotide exchange and auto-phosphorylation in KaiC_RS_, potentially explaining auto-phosphorylation in other KaiBC-based systems lacking a coiled-coil bundle. Notably, the coiled-coil bundle provides additional hexameric stability: the KaiC_RS_ dodecamer is stable for extended periods of time in the presence of only ADP (Extended Data Fig. 3g,h), whereas for KaiC_SE_ oligomers are not observed under these conditions^21^.

**Fig. 2.**
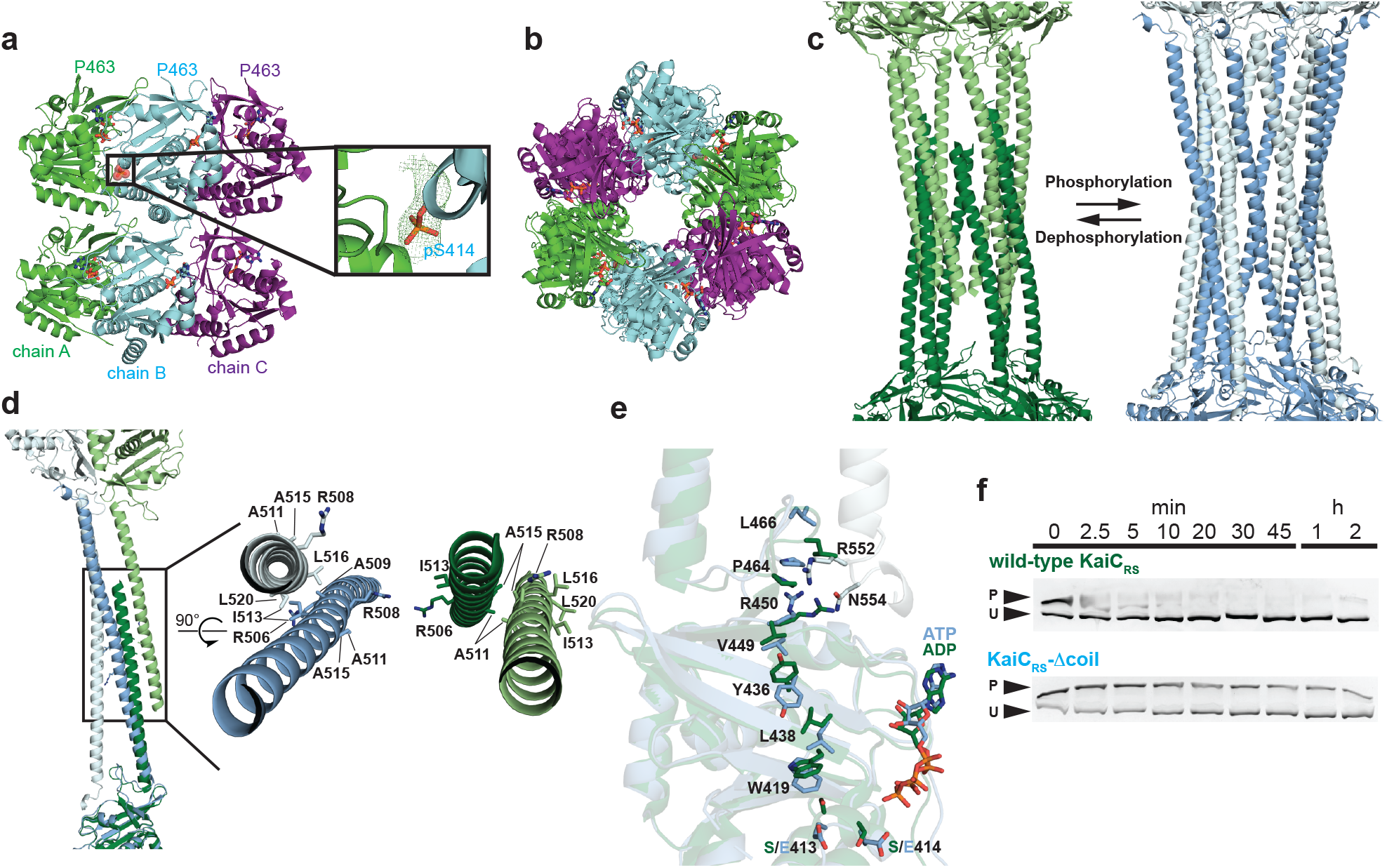
Coiled-coil partner switch coupled to an allosteric network in the CII domain promotes auto-dephosphorylation. **(a)** The X-ray structure of KaiC_RS_-Δcoil was solved in the C222_1_ space group and contained three monomers in the asymmetric unit with ADP present in all active sites. The obtained electron density map allowed for model building up to Pro^463^, indicating that the truncation at position 490 results in enhanced flexibility beyond Pro^463^. Phosphorylation of Ser^414^ was observed in chain B (cyan) as shown by the electron density mFo-DFc polder map (green mesh, 3σ contour level). **(b)** Assembly analysis using the PISA software^37^ revealed a hexamer as the most probable quaternary structure (top view). **(c)** Structural comparison of the coiled-coil domain for unphosphorylated KaiC_RS_ (dark and light green; X-ray structure) and the KaiC_RS_-S413E/S414E phosphomimetic mutant (dark and light blue; cryo-EM structure). (**d**) Overlay of interacting dimers of the structures in (c) using CII domain of chain A as reference (dark shades; bottom). Unphosphorylated KaiC_RS_ (dark green) interacts with the opposite partner on the right (light green), whereas KaiC_RS_-S413E/S414E (dark blue) interacts with the partner on the left (light blue). The hydrophobic packing in the coiled-coil domain is mediated by only the C^β^ atoms of alanine and arginine residues in unphosphorylated KaiC_RS_ but involves the entire side chain of leucine and isoleucine residues in the phosphomimetic structure. (**e**) Allosteric network in the phosphomimetic state (blue) from the coil (light blue) propagating through the KaiC_RS_ CII domain to the active site (dark blue) compared to the unphosphorylated state (dark green). **(f)** Auto-dephosphorylation of KaiC_RS_ and KaiC_RS_-Δcoil over time in the presence of 4 mM ADP at 30 °C. The phosphorylated (P) and unphosphorylated (U) proteins were separated by Zn^2+^ Phos-tag™ SDS-PAGE.

### Long-range allosteric network in KaiC_RS_

The change in phosphorylation state of KaiC has been well established to be the central feature for the circadian rhythm^22,23^. Interestingly, when comparing the unphosphorylated form of full-length KaiC_RS_ (PDB 8dba) and its phosphomimetic mutant (S413E/S414E, PDB xxxx, Extended Data Fig. 4, and Extended Data Table 2) we observe two distinct coiled-coil interactions. Upon phosphorylation, the coiled-coil pairs swap partners by interacting with the other neighboring chain from the opposite hexamer resulting in a register shift that propagates ∼115 Å along the entire coiled-coil (Fig. 2c and Extended Data Fig. 5). In the phosphomimetic state, the register comprises bulkier hydrophobic residues thus resulting in a more stable interaction than for the dephosphorylated form (Fig. 2d and Extended Data Fig. 3g). Furthermore, the C-terminal residues of KaiC_RS_-S413E/S414E interact with the CII domain of the opposite hexamer, whereas the lack of electron density for the last 30 residues in the wild-type structure indicates more flexibility in the dephosphorylated state. Importantly, we discover that these conformational changes in the coiled-coil domain seem to be coupled through a long-range allosteric network to the phosphorylation sites. The rotameric states of residues Ser^413^, Ser^414^, Trp^419^, Val^421^, Tyr^436^, Leu^438^, Val^449^, and Arg^450^ move concertedly and point towards the nucleotide-binding site when the protein is phosphorylated or away in the absence of a phosphate group (Fig. 2e, Extended Data Fig. 5d). We hypothesize that the proximity of the nucleotide to the phosphorylated residue allows for a more efficient phosphoryl transfer and, therefore, determined experimentally the impact of the coiled-coil domain on the auto-dephosphorylation of KaiC_RS_. Indeed, we observe a noticeable effect: wild-type protein dephosphorylates comparatively quickly (*k*_obs_ = 11.5 ± 0.8 h^-1^) in the presence of only ADP, whereas little dephosphorylation was observed for KaiC_RS_-Δcoil (Fig. 2f and Extended Data Fig. 3i) where the allosteric propagation is disrupted (Extended Data Fig. 5d). Consistent with this accelerated dephosphorylation due to the coiled-coil domain, our crystallographic data show a phosphate-group on Ser^414^ for KaiC_RS_-Δcoil but not for the wild-type protein (Fig. 2a and Extended Data Fig. 5d).

### ATP-to-ADP ratio to reset the clock

The surprising result here is that KaiC_RS_ can auto-dephosphorylate on its own despite being constitutively active for phosphorylation due to its extended A-loop conformation. In the canonical *kaiABC* system, the interaction between KaiB and KaiC is required to provide a new binding interface which sequesters KaiA from its “activating” binding site, thereby promoting auto-dephosphorylation at the organism’s optimum temperature^24-26^. The obvious next questions are whether the KaiC_RS_ system can oscillate and secondly if there is a regulatory role for KaiB_RS_ in this process. Comparing the *in vitro* phosphorylation states of KaiC_RS_ in the absence and presence of KaiB_RS_ shows an initial, fast phosphorylation followed by an oscillatory-like pattern in the presence of KaiB_RS_ (hereafter referred to as KaiBC_RS_), whereas KaiC_RS_ alone remains phosphorylated (Fig. 3a,b). Interestingly, the ATP consumption during the reaction with KaiB_RS_ is significantly higher than without (Fig. 3a) and we have seen earlier that KaiC_RS_ will dephosphorylate completely in the presence of only ADP (*cf*. Fig. 2f). These results suggest that the phosphorylation state of KaiC_RS_ and thus the observed oscillatory half-cycle (Fig. 3a,b), is likely related to a change in ATP-to-ADP ratio and we conjectured this could constitute the environmental cue to reset the timer. To test our hypothesis, an ATP-recycling system was added after complete dephosphorylation of KaiBC_RS_ and, as predicted, KaiC_RS_ was able to restart the cycle and phosphorylate again (Extended Data Fig. 6a). Naturally, *in vivo* the ATP-to-ADP ratio will not vary as drastically as in this *in vitro* experiment since nucleotide homeostasis is tightly regulated. To mimic the day- and night-period for *Rhodobacter sphaeroides* we repeated the experiments while keeping the ATP-to-ADP ratio constant (mostly ATP at daytime due to photosynthesis versus 25:75% ATP:ADP during nighttime, respectively)^27^. In the presence of high ATP (i.e., mimicking daytime) KaiC_RS_ remains singly or doubly phosphorylated (Fig. 3c and Extended Data Fig. 6b) irrespective of KaiB_RS_, whereas a constant 25:75% ATP-to-ADP ratio (i.e., mimicking nighttime) results in a much higher fraction of dephosphorylated KaiC_RS_ in the presence of KaiB_RS_ (Fig. 3c). Moreover, when the ATP-to-ADP ratio is flipped to mimic the daytime, KaiC_RS_ is able to phosphorylate again (*cf*. Fig. 3c; around the 28-hour mark). Our data support the notion that the phosphorylation behavior depends strongly on the ATP-to-ADP ratio and, more importantly, demonstrate that the physical binding of KaiB_RS_ results in a higher level of KaiC_RS_ dephosphorylation at nighttime.

**Fig. 3.**
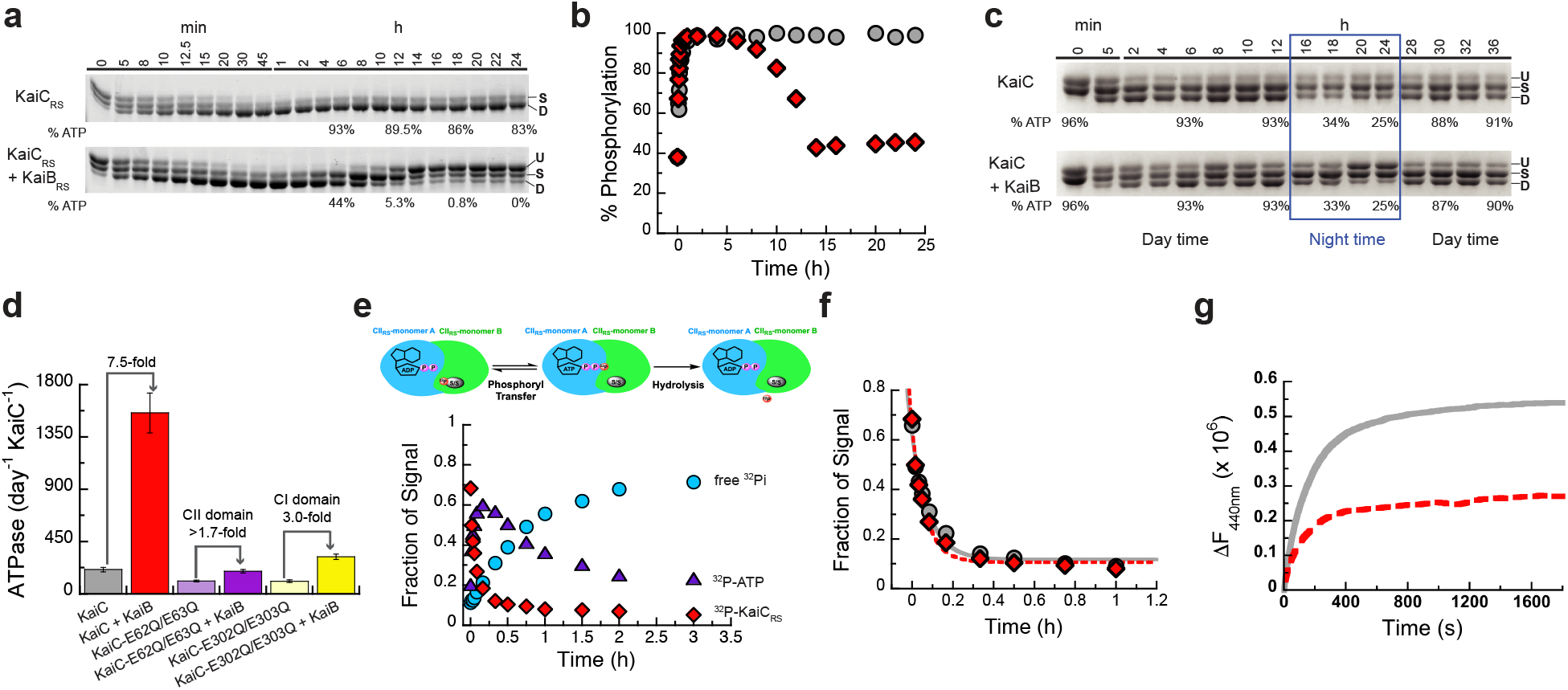
The regulatory role of KaiB_RS_ in the phosphorylation/dephosphorylation cycle of KaiC_RS_. **(a)** SDS-PAGE gel of 3.5 μM KaiC_RS_ and 4 mM ATP in the absence (upper gel) and presence (lower gel) of 3.5 μM KaiB_RS_ at 35 °C with the percentage of ATP indicated at specific time points. U, S, and D represent unphosphorylated, singly, and doubly phosphorylated KaiC_RS_, respectively. **(b)** Phosphorylation (single + double) of KaiC_RS_ during the reaction in the absence (gray circles) or presence (red diamonds) of KaiB_RS_ demonstrates its role in the dephosphorylation of KaiC_RS._ **(c)** Phosphorylation-dephoshorylation cycle of 3.5 μM phosphorylated KaiC_RS_ in the absence and presence of 3.5 μM KaiB_RS_ in a constant ATP-to-ADP ratio of high ATP (4 mM) to mimic daytime and about 25% ATP to mimic the nighttime (exact percentage of ATP indicated at specific time points) at 30 °C. U, S, and D in both panels represent the unphosphorylated, singly phosphorylated (at Ser^413^ or Ser^414^), and doubly phosphorylated state of KaiC_RS_, respectively. These data show that the physical binding of KaiB_RS_ is crucial to dephosphorylate KaiC_RS_ at nighttime where ATP concentrations are still sufficiently high to otherwise keep KaiC_RS_ in the phosphorylated state. **(d)** ATPase activity of wild-type KaiC_RS_ in the absence (gray bar) and presence (red bar) of KaiB_RS_, KaiC_RS_-E62Q/E63Q in the absence (light purple bar) and presence (dark purple bar) of KaiB_RS_, and KaiC_RS_-E302Q/E303Q in the absence (light yellow bar) and presence (dark yellow bar) of KaiB_RS_ at 30 °C. Bar graphs show mean ± S.D from three replicates. **(e)** Time-dependent auto-dephosphorylation of ^32^P-labeled KaiC_RS_ bound with ADP in the presence of 20 μM KaiB_RS_ and 4 mM ADP at 30 °C showing phosphorylated ^32^P-KaiC_RS_ (red diamonds), ^32^P-ATP (purple triangles), and free ^32^Pi (cyan circles). The reaction products were separated by TLC. **(f)** The decay of phosphorylated ^32^P-KaiC_RS_ bound with 4 mM ADP in the absence (gray circles) and presence (red diamonds) of KaiB_RS_ at 30 °C is obtained from autoradiography quantification (see Extended Data Fig. 7). **(g)** The nucleotide exchange of 3.5 μM KaiC_RS_ (gray trace) and 3.5 μM KaiC_RS_ in complex with 30 μM KaiB_RS_ (red dotted trace) in the presence of ATP with mant-ATP. The proteins were incubated at 30 °C for 24 h before the addition of mant-ATP. Representative traces are shown and the fitted parameters (mean ± S.D) were obtained from three replicate measurements.

Next, we investigated the accelerated ATPase activity in KaiC_RS_ upon complex formation. The ATPase activity reported for KaiC_SE_ is very low (∼15 ATP molecules day^-1^ molecule^-1^ KaiC_SE_) and was proposed as a reason for the “slowness”^28^. KaiC_RS_ alone already shows a significantly faster ATPase rate that gets further enhanced by binding of KaiB_RS_ (208 ± 19 and 1,557 ± 172 ATP molecules day^-1^ KaiC_RS_^-1^, respectively; left two bars in Fig. 3d and Extended Data Fig. 6c-g). Furthermore, KaiC_RS_ exhibits no temperature compensation for its ATPase activity (*Q*_10_ ∼1.9; Extended Data Fig. 6c), a feature that is present in KaiC_SE_ and proposed to be a prerequisite for self-sustained rhythms^28^. The deviation from unity for *Q*_10_ is consistent with our earlier observation that the KaiBC_RS_ system is not a true circadian oscillator but rather an hourglass-timer (*cf*. Fig. 3b).

### Regulatory role of KaiB_RS_

The mechanistic details of how KaiB binding in the CI domain allosterically affects the auto-dephosphorylation of KaiC_RS_ in the CII domain remain unclear. Intuitively, there are three plausible scenarios to explain this, namely that KaiB_RS_ binding (i) stimulates the phosphoryl-transfer from pSer back to ADP (Extended Data Fig. 7a), (ii) increases the hydrolysis rate of the active-site ATP (Extended Data Fig. 8a), or (iii) accelerates the nucleotide exchange in the CII domain (Extended Data Fig. 8e). To differentiate between these possibilities, we performed radioactivity experiments to follow nucleotide interconversion, measured ATPase activity for wild-type KaiC_RS_ and mutant forms that are incapable of ATPase activity in the CI or CII domain, and quantified nucleotide-exchange rates by fluorescence using mant-ATP. First, we detected fast, transient ^32^P-ATP formation in our radioactivity experiments when starting from ^32^P-phosphorylated KaiC_RS_ due to its ATP-synthase activity in the CII domain (Fig. 3e and Extended Data Fig. 7b-d). The observed phosphoryl-transfer rate is independent of KaiB_RS_ (*k*_obs_ *=* 12.0 ± 1.7 h^-1^ and 15.4 ± 1.7 h^-1^ in its absence and presence, respectively; Fig. 3f) and agrees well with the rates determined from our gel electrophoresis experiments (11.0 ± 0.8 h^-1^ and 11.5 ± 0.8 h^-1^ with/without KaiB_RS_, respectively; Extended Data Fig. 7e,f). Our experimental data confirm that KaiC_RS_ undergoes dephosphorylation via an ATP-synthase mechanism similarly to what was observed for KaiC_SE29_; KaiB does not expedite the actual phosphoryl-transfer reaction, which is never the rate-limiting step. Since we were unable to stabilize the first phosphorylation site (Ser^414^) in the presence of ADP, the rates reported here correspond exclusively to dephosphorylation of Ser^413^. Secondly, to deconvolute the contributions of the CI and CII domains to the observed ATPase activity, we measured ADP production by KaiC_RS_ mutants that abolish hydrolysis in either the CI (KaiC_RS_-E62Q/E63Q) or CII (KaiC_RS_-E302Q/E303Q) domain. For wild-type KaiC_RS_, the binding of KaiB_RS_ results in a 7.5-fold increase in ATPase activity, and we show that both domains are affected and contribute additively (3-fold for CI and at least 1.7-fold for CII, respectively) to the overall effect (Fig. 3d and Extended Data Fig. 8b-d). Of note, the fold increase in the CII domain represents a lower limit since the KaiC_RS_-E62Q/E63Q mutations interfere with KaiB_RS_ binding as reported before for KaiC_SE30_. Thirdly, our measurements of the nucleotide exchange rate show that it is also unaffected by KaiB_RS_ binding (19.8 ± 1.8 h^-1^ and 18.0 ± 1.5 h^-1^ with/without, respectively; Fig. 3g); since there is no Trp residue near the nucleotide binding site in the CI domain, only the exchange rate in the CII domain could be determined. Interestingly, the change in fluorescence amplitude is much smaller in the presence of KaiB_RS_, demonstrating that even though the binding of KaiB_RS_ does not accelerate nucleotide exchange, it appears to induce a conformational rearrangement in the CII domain especially at higher temperature (Fig. 3g and Extended Data Fig. 8f-h).

### Structure of KaiBC_RS_ complex

To elucidate the structural underpinning of the faster ATPase activity upon KaiB_RS_ binding, we solved the cryo-EM structures of KaiC_RS_ alone (PDB xxxx) and in complex with KaiB_RS_ (PDB xxxx) (Extended Data Table 2). Twelve KaiB_RS_ molecules (monomeric in solution, Extended Data Fig. 9a) bind to the CI domain of the KaiC_RS_-S413E/S414E dodecamer (Fig. 4a-c and Extended Data Fig. 9b). The bound state of KaiB_RS_ adopts the same “fold-switch” conformation as observed for KaiB_TE_^25^ and suggests that this is the canonical binding-competent state (Fig. 4b). Upon binding of KaiB_RS,_ the CI-CI interfaces loosen up (Fig. 4c), which allows for the formation of a tunnel that connects bulk solvent to the position of the hydrolytic water in the active sites (Fig. 4d and Extended Data Fig. 9c). There are other lines of evidence for the weakened interactions within the CI domains. First, KaiB_RS_ binding to either KaiC_RS_-CI domain (Extended Data Fig. 10a) or KaiC_RS_-Δcoil (i.e., missing the C-terminal extensions; Extended Data Fig. 10b) results in disassembly of the hexameric KaiC_RS_ structure into its monomers. In contrast, full-length KaiC_RS_ maintains its oligomeric state upon binding of KaiB_RS_ likely due to the stabilization provided by the coiled-coil interaction. Secondly, a decrease in melting temperature (*T*_M_) of KaiC_RS_ is observed with increasing KaiB_RS_ concentration (Extended Data Fig. 10c). There is no interaction between neighboring KaiB_RS_ molecules within the complex (Extended Data Fig. 9b), suggesting a non-cooperative assembly of KaiB_RS_ to KaiC_RS_ contrary to what is observed for the KaiBC_SE/TE_ complex^31,32^.

**Fig. 4.**
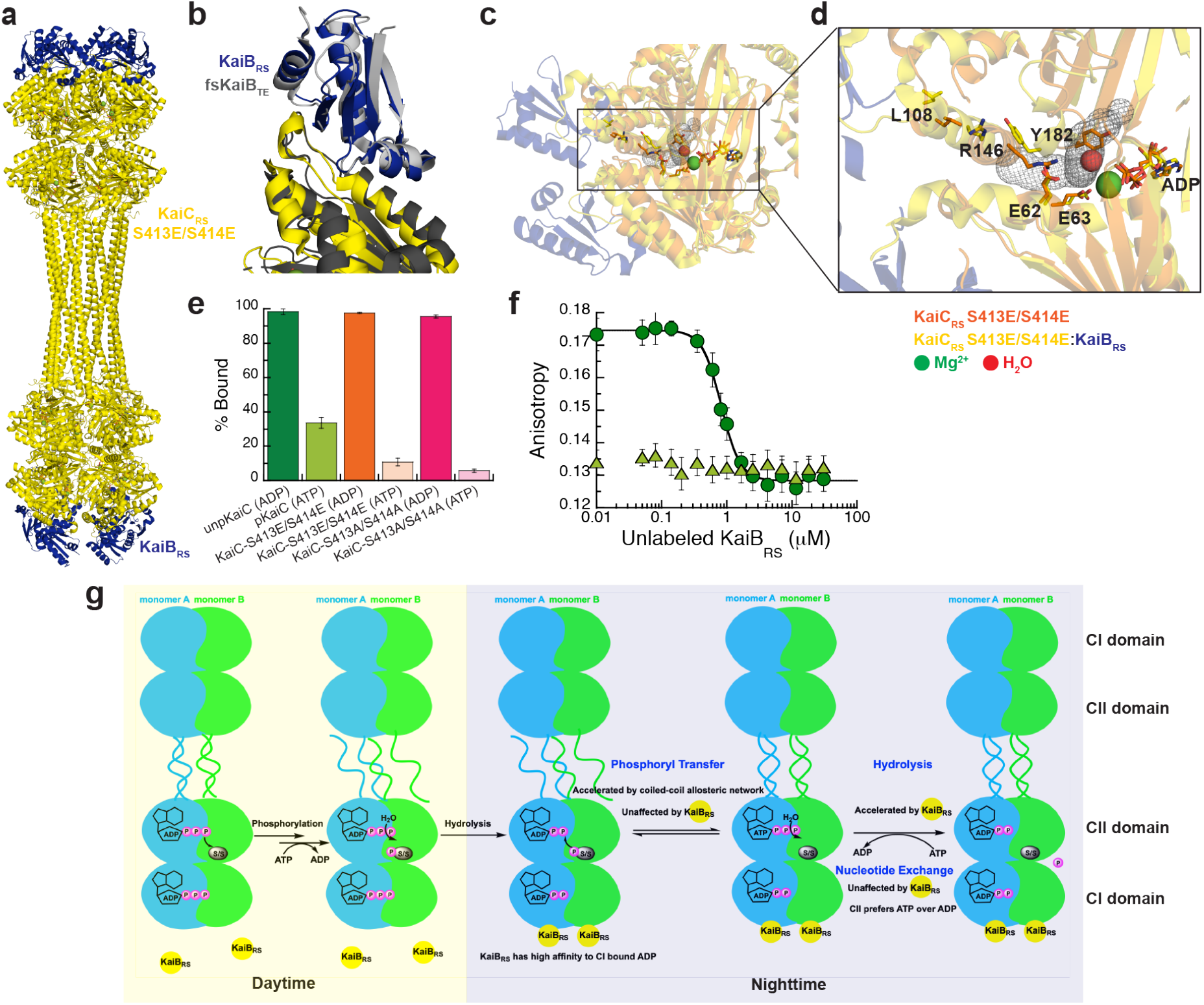
KaiB_RS_ binds to post-hydrolysis state and accelerates the ATPase activity of KaiC_RS_. Cryo-EM structure of KaiC_RS_-S413E/S414E (yellow) in complex with KaiB_RS_ (blue) (PDB xxxx). **(b)** Superposition of KaiC_RS_-S413E/S414E (yellow) bound to KaiB_RS_ (blue) (PDB xxxx) and KaiC_TE_-S413E (dark gray) bound to fsKaiB_TE_ (light gray) (PDB 5jwq^26^). **(c, d)** Binding of KaiB_RS_ (blue) creates a tunnel (gray mesh) that enables water to reach the catalytic position (red sphere) for ATP hydrolysis in the CI domain. **(e)** Binding of wild-type and mutants forms of KaiC_RS_ to His-tagged KaiB_RS_ in the presence of ADP or ATP/recycling system at 25 °C. Bar graphs show mean ± S.D. from three replicates. **(f)** Fluorescence anisotropy of unlabeled KaiB_RS_ competitively displacing KaiB_RS_-6IAF (where 6IAF is the fluorophore) from unphosphorylated KaiC_RS._ in the presence of ADP (dark green circles) and phosphorylated KaiC_RS_ in the presence of the ATP/recycling system (light green triangles) at 30 °C. The average anisotropy and standard error were calculated from ten replicate measurements. **(g)** Schematic diagram of the uncovered mechanism of KaiC_RS_ regulated by coiled-coil interaction and KaiB_RS_ in the CI and CII domain.

Furthermore, we noticed that KaiB-bound structures in phosphomimetic variants of KaiC_RS_ (Fig. 4c,d) and KaiC_SE26_ have ADP bound in their CI domain, demonstrating that the post-hydrolysis state is also the binding-competent state for KaiB_RS_. To test this hypothesis, a His-tagged KaiB_RS_ protein was used in pull-down assays to detect its physical interaction with wild-type and mutant forms of KaiC_RS_ bound with either ADP or ATP. Nearly all KaiB_RS_ is complexed to ADP-bound KaiC_RS_, whereas less than 30% co-elutes in the ATP-bound form regardless of the phosphorylation state (Fig. 4e and Extended Data Fig. 10d,e). The complex formation depends inversely on the ATP-to-ADP ratio (Extended Data Fig. 10f). We performed fluorescence anisotropy competition experiments for a more quantitative description of the binding interaction between KaiC_RS_ and KaiB_RS_: very similar *K*_D_ values were obtained for unphosphorylated, wild-type KaiC_RS_ (Fig. 4f) and its phosphomimetic form (Extended Data Fig. 10g) bound with ADP (0.42 ± 0.03 μM and 0.79 ± 0.06 μM, respectively). No measurable binding curves were obtained for ATP-bound phosphorylated wild-type KaiC_RS_ (Fig. 4f) or KaiC_RS_-S413E/S414E (Extended Data Fig. 10g) with ATP-recycling system, likely due to the small fraction of complex present. Our data show that the post-hydrolysis state in the CI domain is key for KaiB_RS_ binding, whereas the phosphorylation state of KaiC_RS_ has only a marginal effect.

In summary, we unequivocally demonstrate that binding of KaiB_RS_ at the CI domain in the post-hydrolysis state facilitates the hydrolysis of transiently formed ATP after dephosphorylation of KaiC_RS_ in the CII domain (Fig. 4g). Our fluorescence experiments (Fig. 3g and Extended Data Fig. 8f) detect a conformational change in the CII domain upon KaiB_RS_ binding, but we do not observe major structural changes in the cryo-EM structures. Based on the temperature dependence of the fluorescence amplitudes (Extended Data Fig. 8f) we conjecture that the inability to detect conformational differences is likely because of the low temperature. Since the CII domain prefers to bind ATP over ADP (Extended Data Fig. 10h), ATP hydrolysis in the CII domain stimulated by KaiB_RS_ is particularly important to keep KaiC_RS_ in its dephosphorylated state at nighttime, where the exogenous ATP-to-ADP ratio remains sufficiently high to otherwise result in ATP-binding in the CII active site (*cf*. Fig. 3c and Extended Data Fig. 6b).

## Discussion

The KaiBC_RS_ system studied here represents a primordial, hourglass timekeeping machinery and its mechanism provides insight into more evolved circadian oscillators like KaiABC. The dodecameric KaiC_RS_ shows constitutive kinase-activity due to its extended C-terminal tail that forms a coiled-coil bundle with the opposing hexamer and elicits a conformation akin to the exposed A-loop conformation in KaiAC_SE_, and auto-phosphorylation occurs within half an hour. In the KaiABC_SE_ system, the transition from unphosphorylated to doubly phosphorylated KaiC takes place over about twelve hours and the fine-tuning of this first half of the circadian rhythm is accomplished by the emergence of KaiA_SE_ during evolution. The second clock protein, KaiB, binds at the CI domain with the same “fold-switched” state in both systems. The interaction is controlled by the phosphorylation state in the KaiABC_SE_ system, and its sole function is to sequester KaiA_SE_ from the “activating” binding site, whereas KaiB binding directly accelerates the ATPase activity in the KaiBC_RS_ system regardless of the phosphorylation state. The KaiBC_RS_ system requires an environmental switch in ATP-to-ADP concentration to reset the clock and thus follows the day-night schedule when nucleotide concentrations inherently fluctuate in the organism. By contrast, the self-sustained oscillator KaiABC_SE_ remains functional over a wide range of nucleotide concentrations and responds to changes in the ATP-to-ADP ratio by changing its phosphorylation period and amplitude to remain entrained with the day and night cycle^33^.

The novel structural fold of KaiC utilizes the versatile coiled-coil architecture as part of a long-range allosteric network that regulates KaiC_RS_ dephosphorylation. Nature uses conformational changes in coiled-coil domains for a variety of regulatory functions^34^, including the activity of the motor protein dynein in cellular transport of cargo along the actin filament^12^. A similar register shift, although in a coiled-coil interaction formed by only two helices is used in dynein motility. Given that this simple heptad repeat sequence emerged multiple times and is found throughout all kingdoms of life^35^ it is an example of convergent evolution.

## Data availability

Structure factors and refined models are deposited in the Protein Data Bank (PDB) under accession codes 8dba (wild-type KaiC_RS_, x-ray), 8db3 (KaiC_RS_-Δcoil, x-ray), xxxx (KaiC_RS_-S413E/S414E, cryo-EM), and xxxx (KaiC_RS_-S413E/S414E:KaiB_RS_, cryo-EM), respectively. Cryo-EM maps are deposited in the Electron Microscopy Data Bank (EMDB) under accession codes xxxx (KaiC_RS_-S413E/S414E) and xxxx (KaiC_RS_-S413E/S414E:KaiB_RS_), respectively.

## Acknowledgements

D.K. and N.G. are supported by the Howard Hughes Medical Institute (HHMI). We would like to thank Mike Rigney for assistance with negative-stain data collection at the Brandeis University Electron Microscopy Facility, and Zhiheng Yu and the staff of the Janelia Research Campus cryo-EM facility for advice and assistance with data collection. The Berkeley Center for Structural Biology is supported in part by HHMI. Beamline 8.2.1 of the Advanced Light Source, a U.S. DOE Office of Science User Facility under Contract No. DE-AC02-05CH11231, is supported in part by the ALS-ENABLE program funded by the National Institutes of Health, National Institute of General Medical Sciences, grant P30 GM124169-01. Mass spectral data were obtained at the University of Massachusetts Mass Spectrometry Core Facility, RRID:SCR_019063.

## Author contributions

W.P., R.A.P.P., and D.K. conceived the project and designed experiments. W.P. performed and analyzed all biochemical data. W.P. and R.A.P.P. set up the crystal trays and R.A.P.P. collected and analyzed the X-ray crystallographic data. W.P. prepared the samples for the cryo-EM studies and collected negative stain images to screen for optimal sample conditions; T.G. collected and processed all cryo-EM data, and reconstructed the cryo-EM maps under supervision of N.G.; R.A.P.P. built and interpreted the structural models. W.P. and N.B. performed and analyzed experiments with radioactively labeled KaiC. M.H. built the KaiC phylogeny. W.P., R.O., and D.K. wrote the paper; and all authors commented on the manuscript and contributed to data interpretation.

## Competing interests

D.K. is co-founder of Relay Therapeutics and MOMA Therapeutics. The remaining authors declare no competing interests.

**Extended Data Fig. 1.**
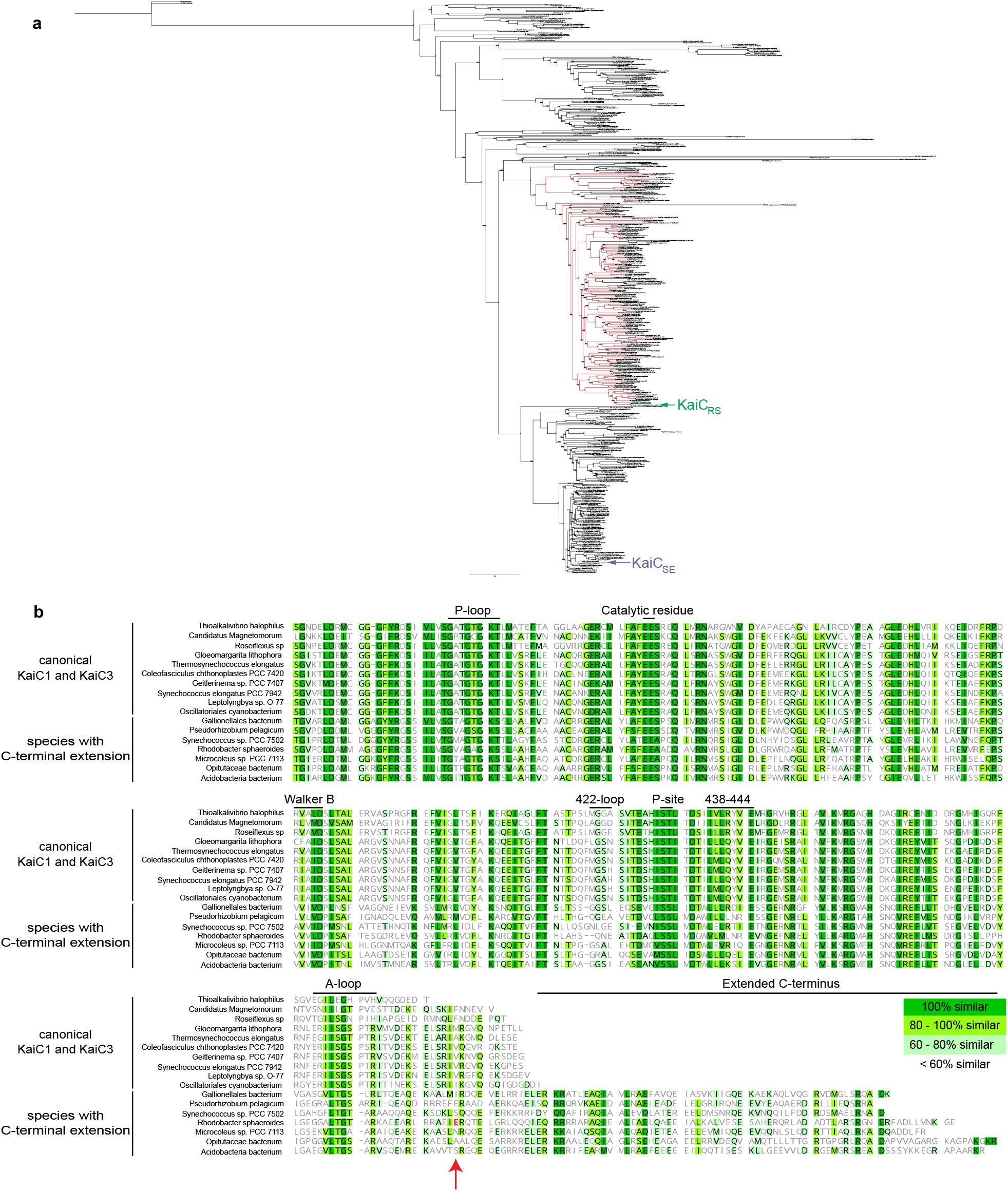
Evolution of *kaiC* and sequence alignment of *kaiC* subgroups. **(a)** Phylogenetic tree of *kaiC*homologs, where *kaiC* genes that have an approximately 50 amino acids C-terminal extension are labeled in red *Rhodobacter sphaeroides* strain KD131 studied here and *Synechococcus elongatus* PCC 7942 (widely studied in the literature) are highlighted in green and pink, respectively. The accession code and organism are shown at the tip of the branches, the numbers at each node represent the aBayes bootstrap values^38^, and the legend for branch length is shown. **(b)** A sequence alignment of the CII domain of the *kaiC* subgroups annotated with its sequence similarity. Residue Glu^490^, the position where the stop codon was introduced in the truncated KaiC_RS_-Δcoil construct, is shown in red and marked with an arrow.

**Extended Data Fig. 2.**
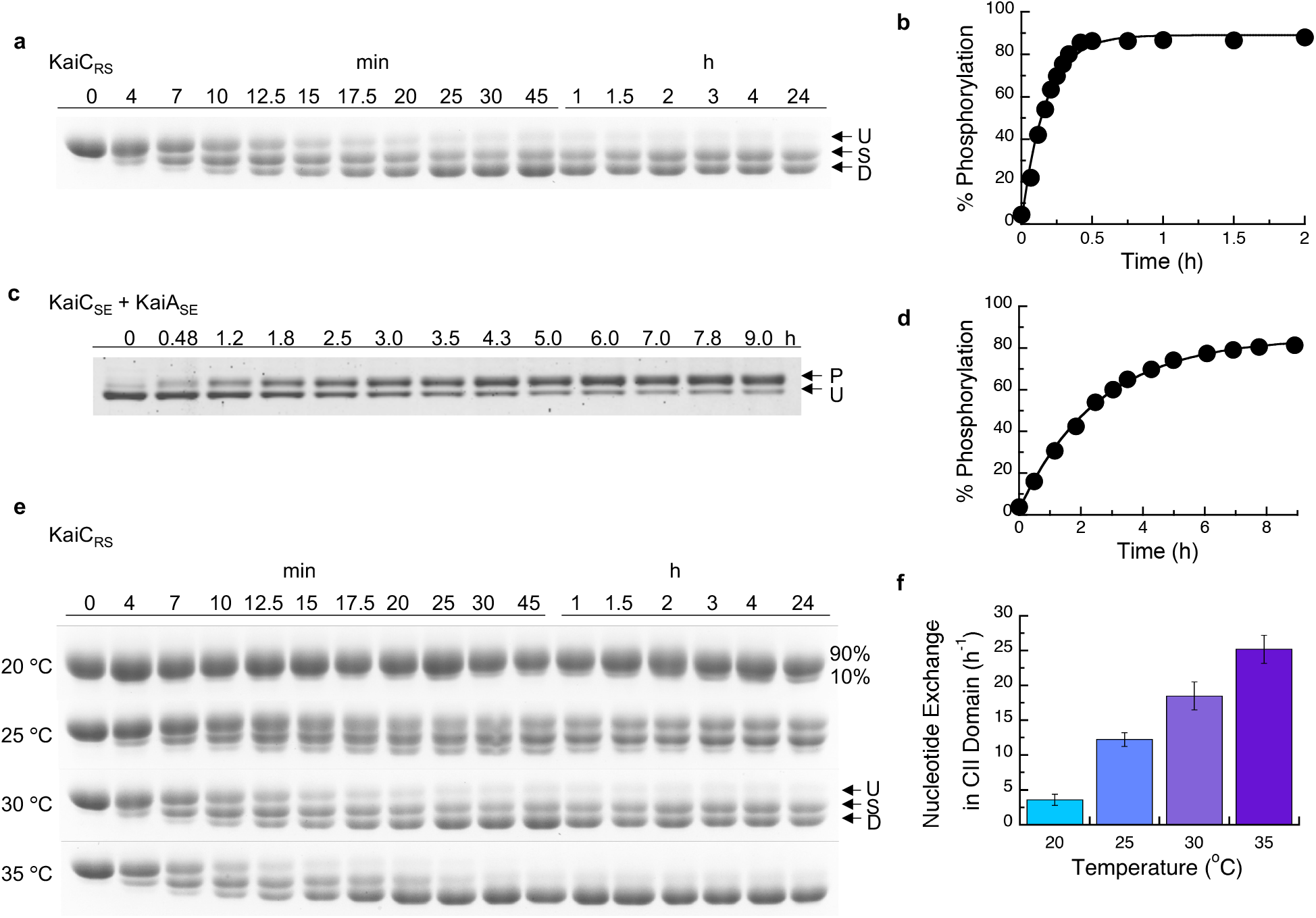
Auto-phosphorylation and nucleotide exchange rates of KaiC. **(a)** 10% SDS-PAGE gel of 3.5 μM KaiC_RS_ in the presence of 4 mM ATP and using an ATP-recycling system at 30 °C. U, S, and D represent unphosphorylated, singly, and doubly phosphorylated KaiC_RS_, respectively. **(b)** Densitometric analysis of auto-phosphorylation (single + double phosphorylation) from panel (a) over time yields a rate of 6.5 ± 1.0 h^-1^. **(c)** 6.5% SDS-PAGE gel of 3.5 μM KaiC_SE_ in the presence of 1.2 μM KaiA_SE_ and 4 mM ATP at 30 °C. U and P represent unphosphorylated and phosphorylated KaiC_SE_, respectively. **(d)** Densitometric analysis of auto-phosphorylation of KaiC_SE_ activated by KaiA_SE_ (panel (c)) shows a rate of 0.40 ± 0.02 h^-1^ and is substantially slower than for KaiC_RS_. The standard deviation for parameters in (b) and (d) were obtained from data fitting. **(e)** 10% SDS-PAGE gels for experiments with 3.5 μM KaiC_RS_ in the presence of 4 mM ATP and using an ATP-recycling system between 20 and 35 °C show that the level of phosphorylation increases with temperature. U, S, and D represent unphosphorylated, singly, and doubly phosphorylated KaiC_RS_, respectively. **(f)** Bar graphs indicating the nucleotide exchange rate in the CII domain of KaiC_RS_ incubated with 50 μM ATP in the presence of an ATP-recycling system, and then mixed with 250 μM mant-ATP. An increase in fluorescence intensity at 440 nm was recorded and the single-exponential time traces were fitted to obtain the exchange rate constants: 3.6 ± 0.8 h^-1^ (20 °C), 12.2 ± 1.0 h^-1^ (25 °C), 18.5 ± 1.5 h^-1^ (30 °C), and 25.2 ± 0.2 h^-1^ (35 °C). Experiments were performed in triplicate with error bars representing SD.

**Extended Data Fig. 3.**
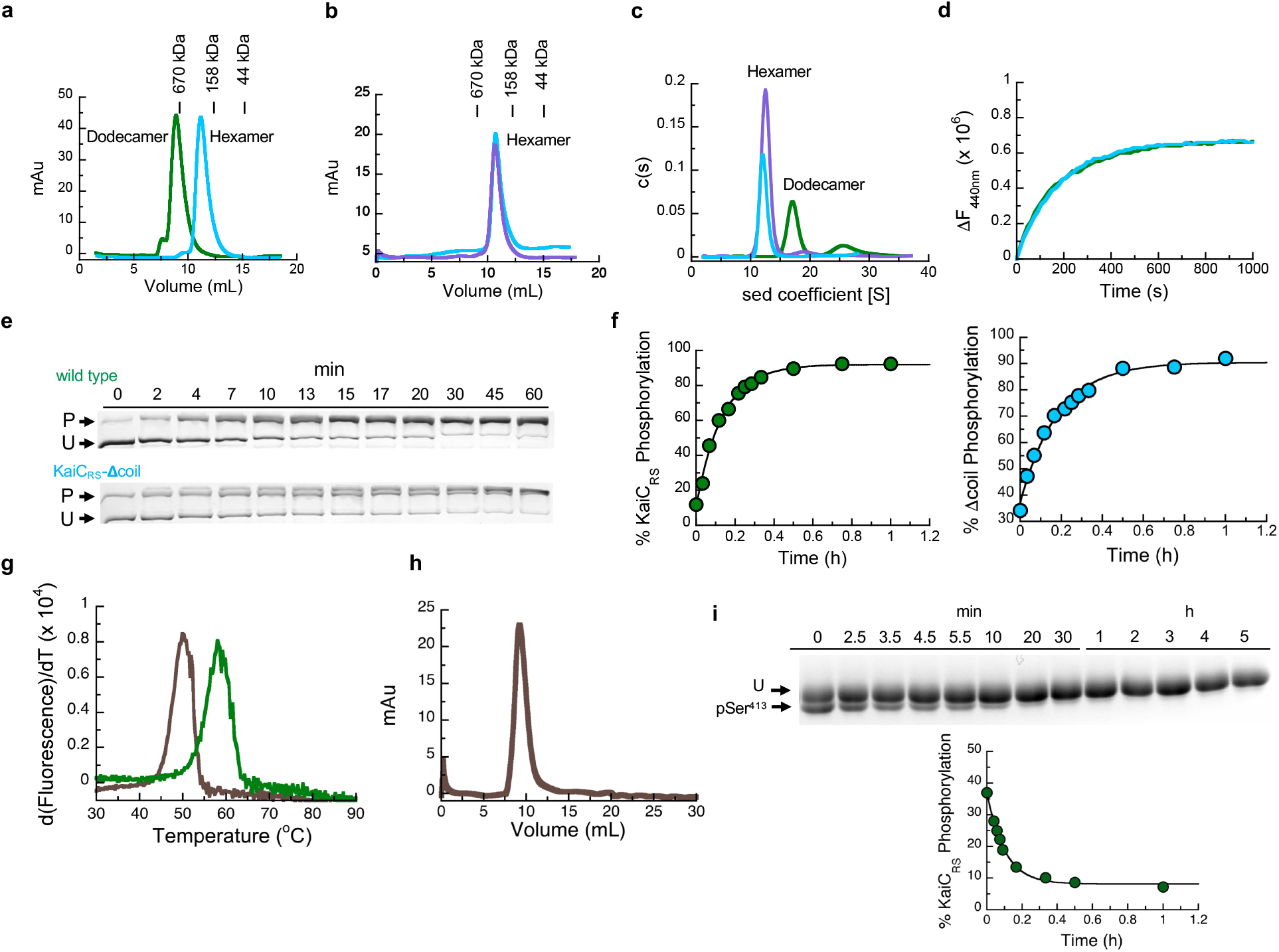
Oligomeric states of KaiC_RS_ and effect of coiled-coil domain on rates of nucleotide exchange and auto-phosphorylation. **(a)** Oligomerization analysis of KaiC_RS_ (dodecamer, green line) and truncated KaiC_RS_-Δcoil (hexamer, cyan line) by analytical gel-filtration chromatography. The protein size markers are indicated at the top. **(b)** Comparison of the elution profiles of KaiC_RS_-Δcoil (cyan line) and KaiC_SE_ (purple line) from size-exclusion chromatography shows a hexameric state for both KaiC_RS_-Δcoil and KaiC_SE_. **(c)** Oligomeric states of KaiC_RS_ (dodecamer, green line), KaiC_RS_-Δcoil (hexamer, cyan line), and KaiC_SE_ (hexamer, purple line) were also measured by analytical ultracentrifugation (sedimentation velocity at 30,000 rpm and 20 °C) and the results agree with the data shown in panels (d) and (e). The graph in panel (f) represents the sedimentation coefficient distribution [c(s)]. **(d)** The change in fluorescence at 440 nm (ΔF_440nm_) represents the nucleotide exchange between ATP and mant-ATP at 30 °C for KaiC_RS_ (green trace, 18.0 ± 1.5 h^-1^) and KaiC_RS_-Δcoil (cyan trace, 19.1 ± 0.8 h^-1^). Representative traces are shown and the fitted parameters (mean ± S.D) were obtained from three replicate measurements. **(e)** Zn^2+^ Phos-tag™ SDS-PAGE gel shows the level of phosphorylation over time of KaiC_RS_ (upper gel) and KaiC_RS_-Δcoil (lower gel) at 35 °C. P and U represent phosphorylated and unphosphorylated protein, respectively. **(f)** Phosphorylation level over time of KaiC_RS_ (green circles, 7.4 ± 0.3 h^-1^) and KaiC_RS_-Δcoil (cyan circles, 5.5 ± 0.4 h^-1^) analyzed by densitometric analysis of Zn^2+^ Phos-tag™ SDS-PAGE gel in (b). **(g)** First derivative of thermal-stability curves measured for unphosphorylated KaiC_RS_ bound with ADP (brown line) and phosphorylated KaiC_RS_ bound with ATP (green line). The extracted temperatures of denaturation are 50 °C (unphosphorylated KaiC_RS_ in the presence of 1 mM ADP) and 58 °C (phosphorylated KaiC_RS_ in the presence of 1 mM ATP), respectively. **(h)** Dodecameric state of unphosphorylated KaiC_RS_ (40 μM) bound with ADP measured by size-exclusion chromatography. **(i)** SDS-PAGE gel shows dephosphorylation of Ser^413^ over time at 30 °C in the presence of 4 mM ADP (U and pSer^413^ represent unphosphorylated and Ser^413^-phosphorylated KaiC_RS_, respectively) with the corresponding kinetics shown in the lower panel (confirmed by MS/MS) with a rate constant of 11.5 ± 0.8 h^-1^. This result suggests that the coiled-coil domain promotes KaiC_RS_ dephosphorylation. The standard deviation for parameters in (f) and (i) were obtained from data fitting.

**Extended Data Fig. 4.**
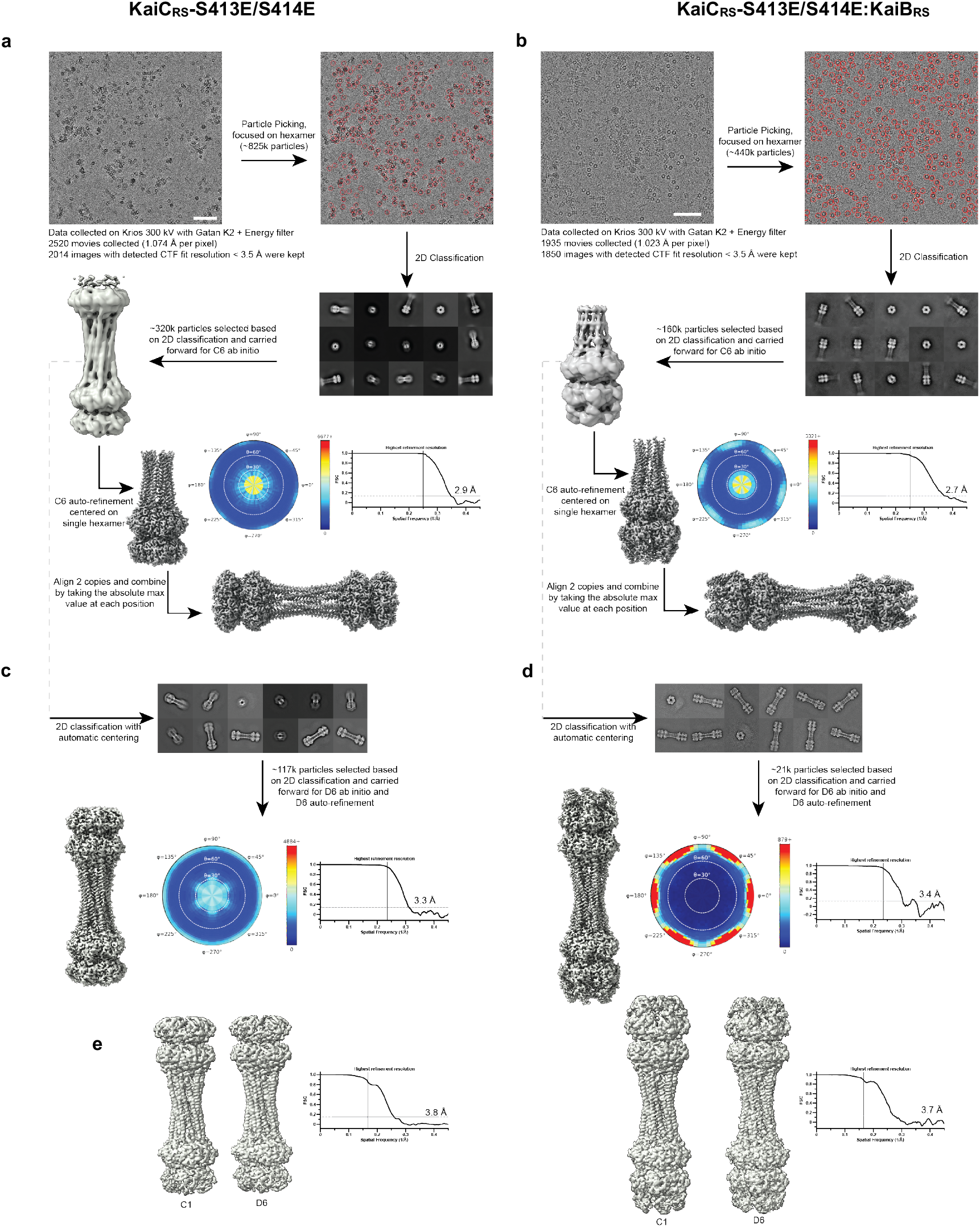
Graphical description of the cryo-EM processing workflow and validation of the final dodecamer structures. The workflow (see description in Methods section) demonstrates a typical image (scale bar: 60nm) and representative good class averages. The ab initio and final reconstructions are shown. Shown alongside the final reconstruction is the angular plot demonstrating the distribution of particle views and the Fourier shell correlation curve used for the global resolution estimation **(a)** KaiC_RS_-S413E/S414E alone and **(b)** KaiC_RS_-S413E/S414E:KaiB_RS_ complex. To validate the final combined dodecamer structures, the data were reprocessed for the full dodecamer. The figure shows representative good class averages, the final reconstruction, angular distribution and Fourier shell correlation curve for the KaiC_RS_-S413E/S414E alone and **(d)** KaiC_RS_-S413E/S414E:KaiB_RS_ complex dodecamers. **(e)** Comparison of the C1 and D6 reconstructions of KaiC_RS_-S413E/S414E alone and KaiC_RS_-S413E/S414E:KaiB_RS_, and Fourier shell correlation curves for the C1 reconstructions. The C1/D6 comparisons do not reveal discernable differences, suggesting that these complexes have D6 symmetry.

**Extended Data Fig. 5.**
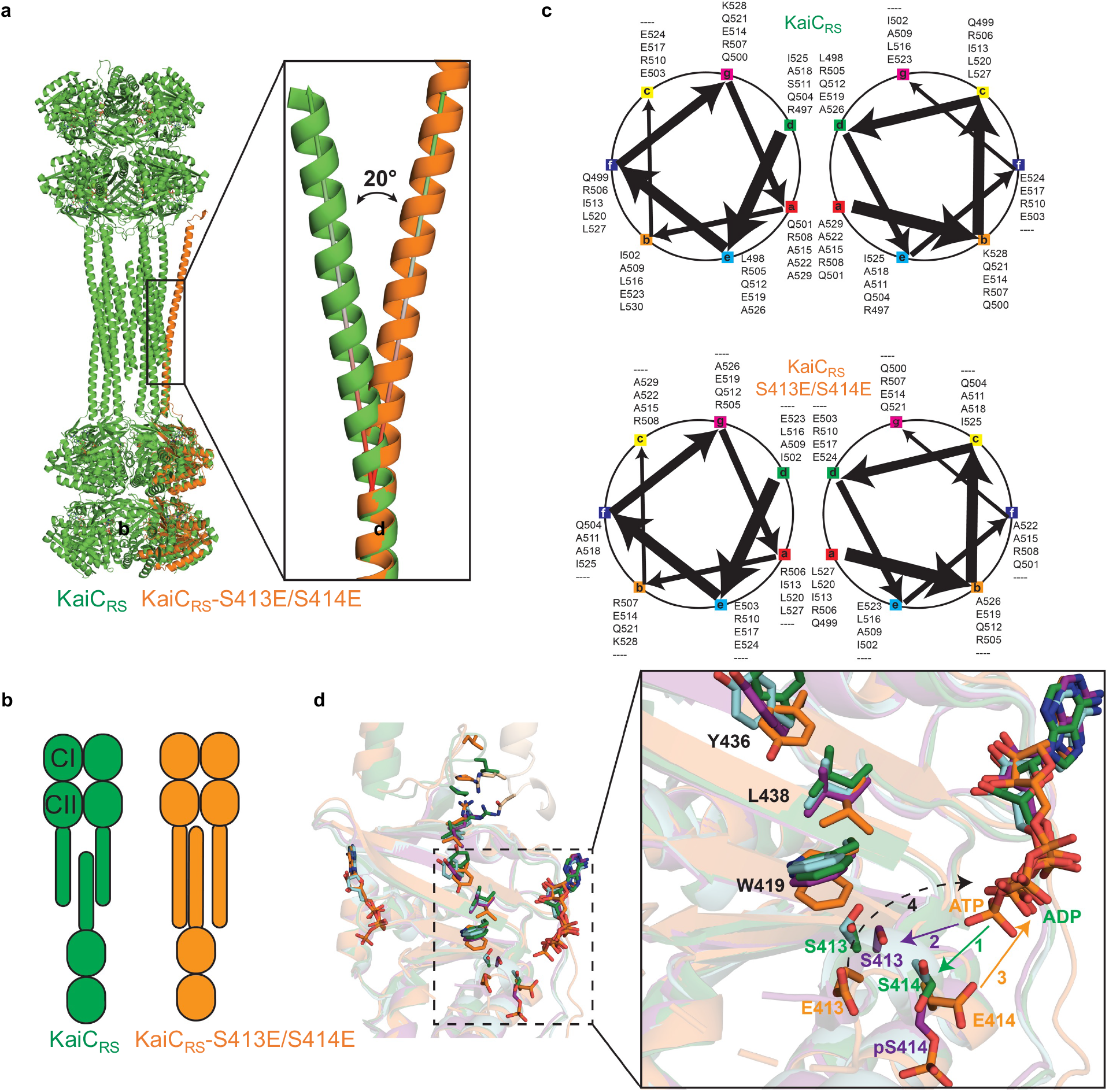
Correlation between the coiled-coil register shift and phosphorylation, and model for consecutive phosphorylation/dephosphorylation events in CII domain of KaiC_RS_. **(a)** Structural comparison between KaiC_RS_ (green) and KaiC_RS_-S413E/S414E (orange, single-chain for clarity) reveals that the coiled-coil in the phosphomimetic structure points outwards, with an angle of about 20° relative to the KaiC_RS_ coiled-coil. **(b)** The conformational change in the coiled-coil domain affects the dimer interface due to partner swaps with the opposite hexamer (see also Fig. 2). From an “outside perspective”: the C-terminal helix in KaiC_RS_ interacts with the right chain from the opposite hexamer, whereas in KaiC_RS_-S413E/S414E the interaction is with the chain on the left. **(c)** Coiled-coil diagrams describe the heptad register shift that accompanies this structural rearrangement. **(d)** Based on the overlay of our structures, we propose the following model for the phosphorylation/ dephosphorylation events. First, the phosphorylation cycle starts with the transfer of the *γ*-phosphate of ATP to the hydroxyl group of Ser^414^ (1; green arrow) in unphosphorylated KaiC_RS_ (green) or KaiC_RS_-Δcoil (cyan). Secondly, pSer^414^ of KaiC_RS_-Δcoil (purple, singly phosphorylated) moves away from the active site placing the hydroxyl group of Ser^413^ closer to the -phosphate of ATP for the second phosphorylation (2; purple arrow). Thirdly, the doubly phosphomimetic state (KaiC_RS_-S413E/S414E, orange) reveals that the phosphoryl group of pSer^414^ moves back towards the active site for dephosphorylation (3; orange arrow). Lastly, we hypothesize that the indole group of Trp^419^ “pushes” pSer^413^ into the active site for the second dephosphorylation event (dashed arrow), in agreement with the slower dephosphorylation rate observed in the KaiC_RS_-Δcoil construct (*cf*. Fig. 2d).

**Extended Data Fig. 6.**
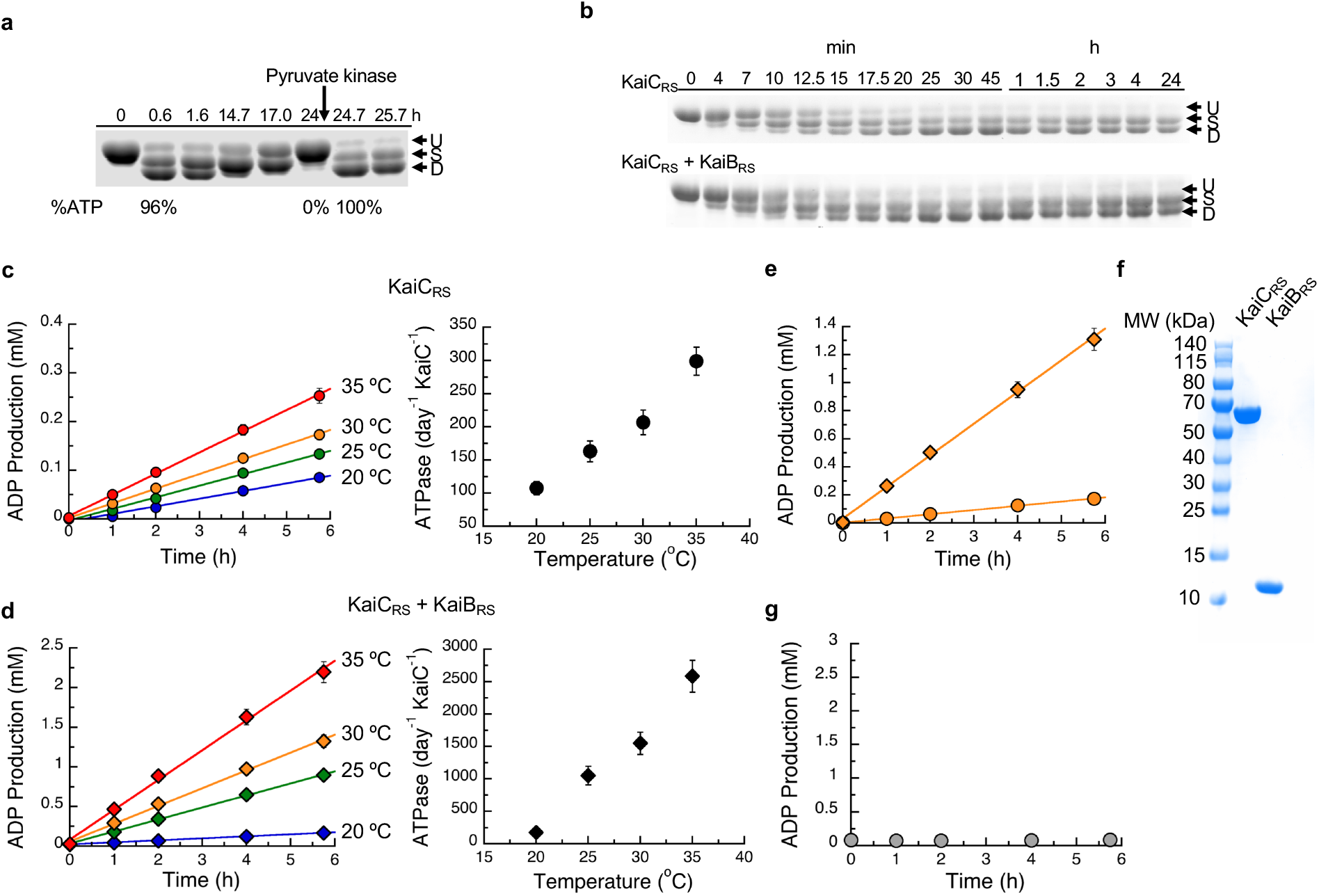
Effect of ATP-to-ADP ratio on KaiC_RS_ auto-dephosphorylation and the dependence of temperature and KaiB_RS_ binding on the ATPase activityof KaiC_RS_. **(a)** 10% SDS-PAGE gel of 3.5 μM KaiCRS and 3.5 μM KaiBRS in the presence of 4 mM ATP and 10 mM 2-phosphoenolpyruvate at 30 °C shows that the auto-phosphorylation cycle restarts upon regeneration of ATP by the addition of 2 U/mL pyruvate kinase at the 24-hour time mark. **(b)** 10% SDS-PAGE gel of 3.5 μM KaiC_RS_ without (upper panel) and with (lower panel) 3.5 μM KaiB_RS_ in the presence of 4 mM ATP with an ATP-recycling system added from the beginning showing that under these conditions KaiB does not accelerate dephosphorylation. **(c)** Representative curves for ADP production of KaiC_RS_ (3.5 μM) alone and **(d)** in the presence of KaiB_RS_ (3.5 μM) in 4 mM ATP measured by HPLC. The data were analyzed as described in the Methods section and result in ATPase activities of 108 ± 10 day^-1^ KaiC^-1^ (with KaiB_RS_ = 176 ± 29 day^-1^ KaiC^-1^) at 20 °C, 163 ± 16 day^-1^ KaiC^-1^ (with KaiB_RS_ = 1052 ± 143 day^-1^ KaiC^-1^) at 25 °C, 208 ± 19 day^-1^ KaiC^-1^ (with KaiB_RS_ = 1557 ± 172 day^-1^ KaiC^-1^) at 30 °C, and 300 ± 21 day^-1^ KaiC^-1^ (with KaiB_RS_ = 2584 ± 245 day^-1^ KaiC^-1^) at 35 °C. The temperature coefficient, *Q*_10_, was calculated using the data obtained at 25 °C and 35 °C and yields a value of ∼ 1.9. The standard deviations of ATPase activity at each temperature were obtained from three replicate measurements. **(e)** The comparison of ADP production of KaiC_RS_ in the absence (orange circles) and presence (orange diamonds) of KaiB_RS_ at 30 °C indicate a 7.5-fold increase in ATPase activity for the complex. The binding of KaiB_RS_ accelerates the ATPase activity of KaiC_RS_ in both the CI and CII domains (see also Extended Data Fig. 8b, c). **(f)** The SDS-PAGE gel of KaiC_RS_ (10 μg) and KaiB_RS_ (10 μg) shows that both proteins were purified to homogeneity and the measured ATPase activity is, therefore, not due to impurities. **(g)** ADP production of KaiB_RS_ in 4 mM ATP at 30 °C shows, as expected, no ATPase activity for KaiB_RS_ alone and confirms the increase in ATPase activity shown in panel (d) is due to complex formation.

**Extended Data Fig. 7.**
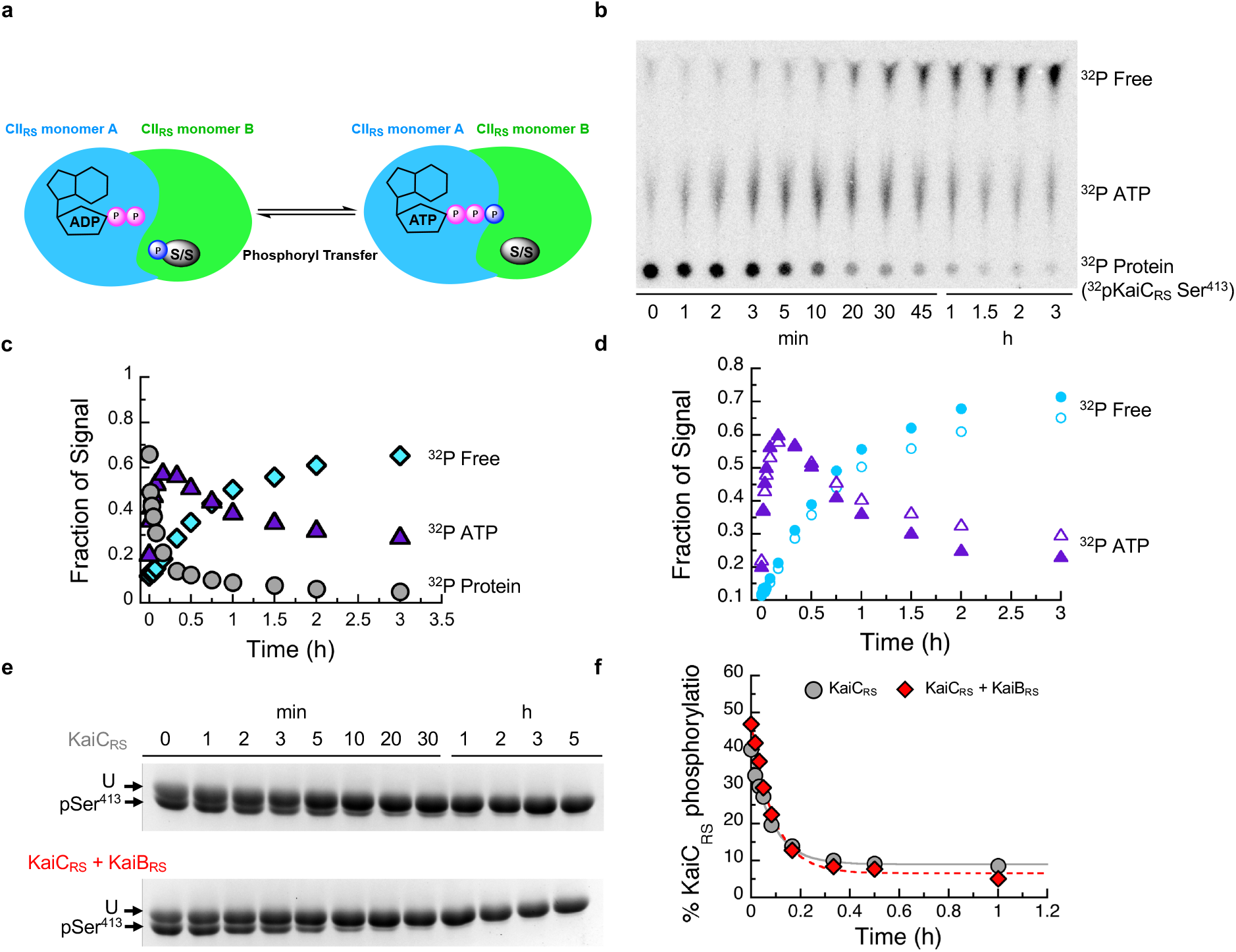
Dephosphorylation of KaiC_RS_ occurs via an ATP-synthase mechanism and the phosphoryl-transfer step is unaffected by the binding of KaiB_RS_. **(a)** Possible mechanisms how KaiB_RS_ could accelerate KaiC_RS_ dephosphorylation at nighttime. Binding of KaiB_RS_ on the CI_RS_ domain directly accelerates the phosphoryl transfer from pSer to bound ADP to generate transiently bound ATP. The cartoon represents the interface between two monomers in the CII_RS_ domain. **(b)** Autoradiograph of separation of ^32^P-KaiC_RS_ at Ser^413^, transiently formed ^32^P-ATP, and free ^32^Pi via thin-layer chromatography (TLC) with 4 mM ADP at 30 °C, with the corresponding kinetics shown in **(c)** where gray circle, purple triangle, and cyan diamonds represent the relative concentrations of phosphorylated ^32^P-KaiC_RS_, ^32^P-ATP, and free ^32^Pi, respectively. **(d)** Comparison of transient ^32^P-ATP formation and decay in the absence (open triangle) and presence (solid triangle) of KaiB_RS_ and free ^32^P formation in the absence (open circles) and presence of KaiB_RS_ (solid circles). Faster decay of transient ^32^P-ATP together with higher free ^32^P production in the presence of KaiB_RS_ indicated that KaiB_RS_ accelerates hydrolysis in KaiC_RS_. **(e)** SDS-PAGE gel (10%) of dephosphorylation of phosphorylated 3.5 μM KaiC_RS_ at Ser^413^ without (upper gel) and with (lower gel) 3.5 μM KaiB_RS_ in the presence of 4 mM ADP at 30 °C. **(f)** Densitometric analysis of data in panel (e) shows the decay of total KaiC_RS_ phosphorylation in the absence (gray circles) and presence (red diamonds) of KaiB_RS_ and yields rates of 11.5 ± 0.8 h^-1^ and 11.0 ± 0.8 h^-1^, respectively. This result indicates that binding of KaiB_RS_ does not accelerate the phosphoryl-transfer step in KaiC_RS_.

**Extended Data Fig. 8.**
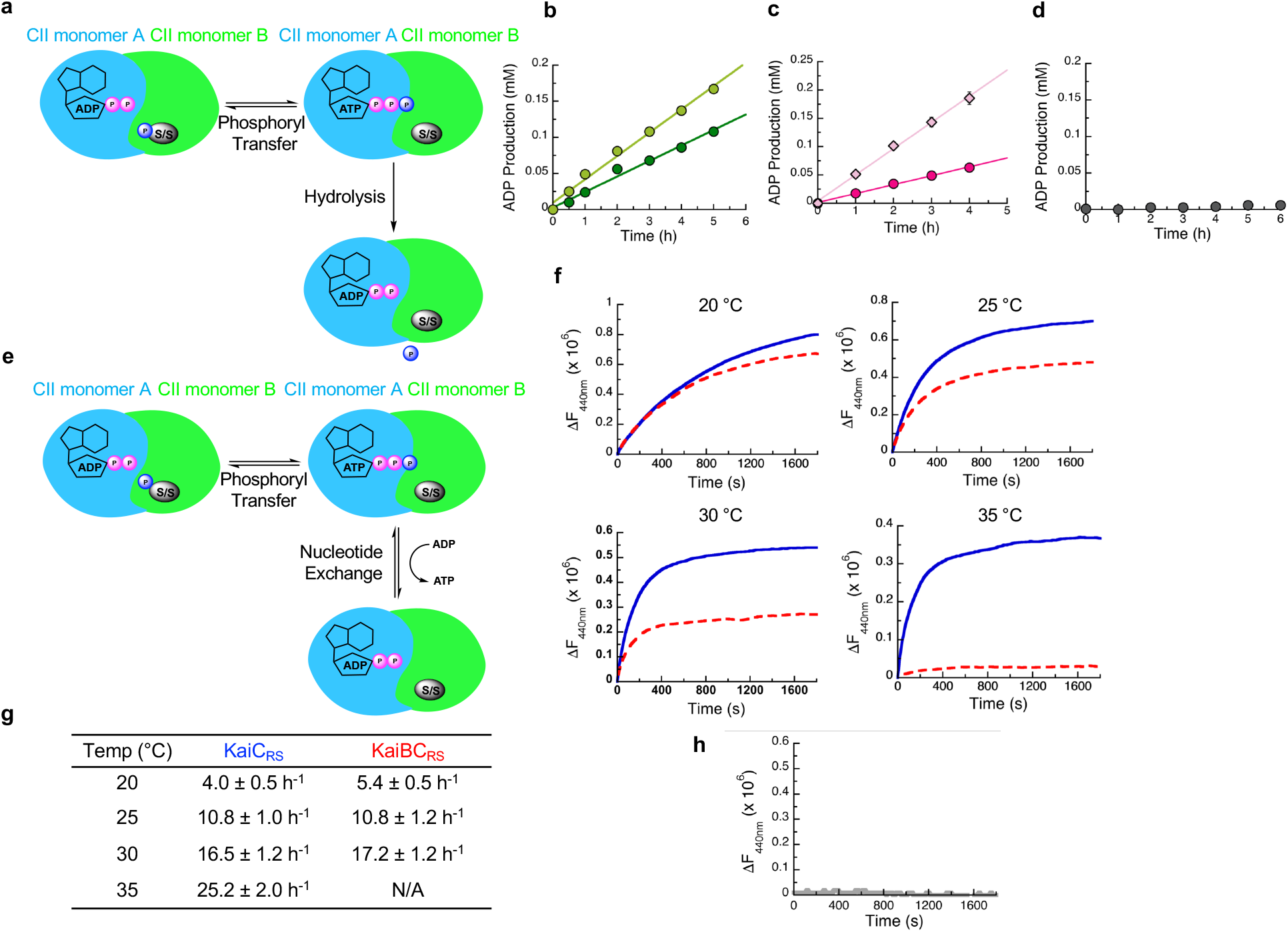
Effect of KaiB_RS_ binding on ATPase activity and nucleotide exchange in the CII domain of KaiC_RS_. **(a)** Second possible mechanism to explain how KaiB_RS_ accelerates KaiC_RS_ dephosphorylation at nighttime: binding of KaiB_RS_ to the CI_RS_ domain could increase the hydrolysis rate in the CII_RS_ domain and, thereby, prevent the phosphoryl transfer back from transiently formed or external ATP back to serine residues. **(b)** ADP production of phosphorylated KaiC_RS_ with catalytic mutations in the CI domain (KaiC_RS_-E62Q/E63Q, 3.5 μM) in the absence (dark green circles) and presence (light green circles) of 3.5 μM KaiB_RS_ at 30 °C with 4 mM ATP was quantified using HPLC. From these data an ATPase activity in the CII domain of 112± 8 day^-1^ KaiC^-1^ and 195 ± 16 day^-1^ KaiC^-1^ in the presence of KaiB_RS_ was determined. **(c)** ADP production measured by HPLC as in panel (b) of KaiC_RS_ but with catalytic mutations in the CII domain (KaiC_RS_-E302Q/E303Q, 3.5 μM) in the absence (dark pink circles) and presence (light pink diamonds) of 3.5 μM KaiB_RS_. The corresponding ATPase activities in the CI domain are 110 ± 12 day^-1^ KaiC^-1^ in the absence and 320± 22 day^-1^ KaiC^-1^ in the presence of KaiB_RS_. **(d)** ADP production of KaiCI_RS_-E62Q/E63Q (construct of only CI domain with catalytic mutations) in 4 mM ATP at 30 °C shows no ATPase activity indicating that Glu^62^ and Glu^63^ are the only two residues that are responsible for ATPase activity in CI domain of KaiC_RS_ and confirms the ATPase activity shown in panel C is due to ATPase activity in CII domain of KaiC_RS_. **(e)** Third possible mechanism to explain how KaiB_RS_ accelerates KaiC_RS_ dephosphorylation at nighttime: binding of KaiB_RS_ to the CI_RS_ domain could promote faster nucleotide exchange in the CII_RS_ domain to displace transient ATP by ADP. **(f)** Time course of fluorescence intensity at 440 nm due to binding of mant-ATP to KaiC_RS_-S413E bound with ATP in the absence (solid blue trace) and presence (red dotted trace) of KaiB_RS_. KaiC_RS_-S413E (3.5 μM) was pre-incubated with 3.5 μM KaiB_RS_ for 16 h at 20, 25, 30, and 35 °C in the presence of 50 μM ATP and an ATP-recycling system and then mixed with 250 μM mant-ATP. The observed exchange rates at each temperature are listed in the table **(g). (h)** Nucleotide exchange of KaiCI_RS_ (i.e., only CI_RS_ domain) cannot be measured since there is no tryptophan residue in close proximity of the nucleotide binding site. In summary, KaiB_RS_ accelerates KaiC_RS_ dephosphorylation by increasing the hydrolysis rate in the CI and CII domains and does not affect the nucleotide exchange rate. Representative traces are shown in (f) and (h) and the fitted parameters (g; mean ± S.D) were obtained from three replicate measurements.

**Extended Data Fig. 9.**
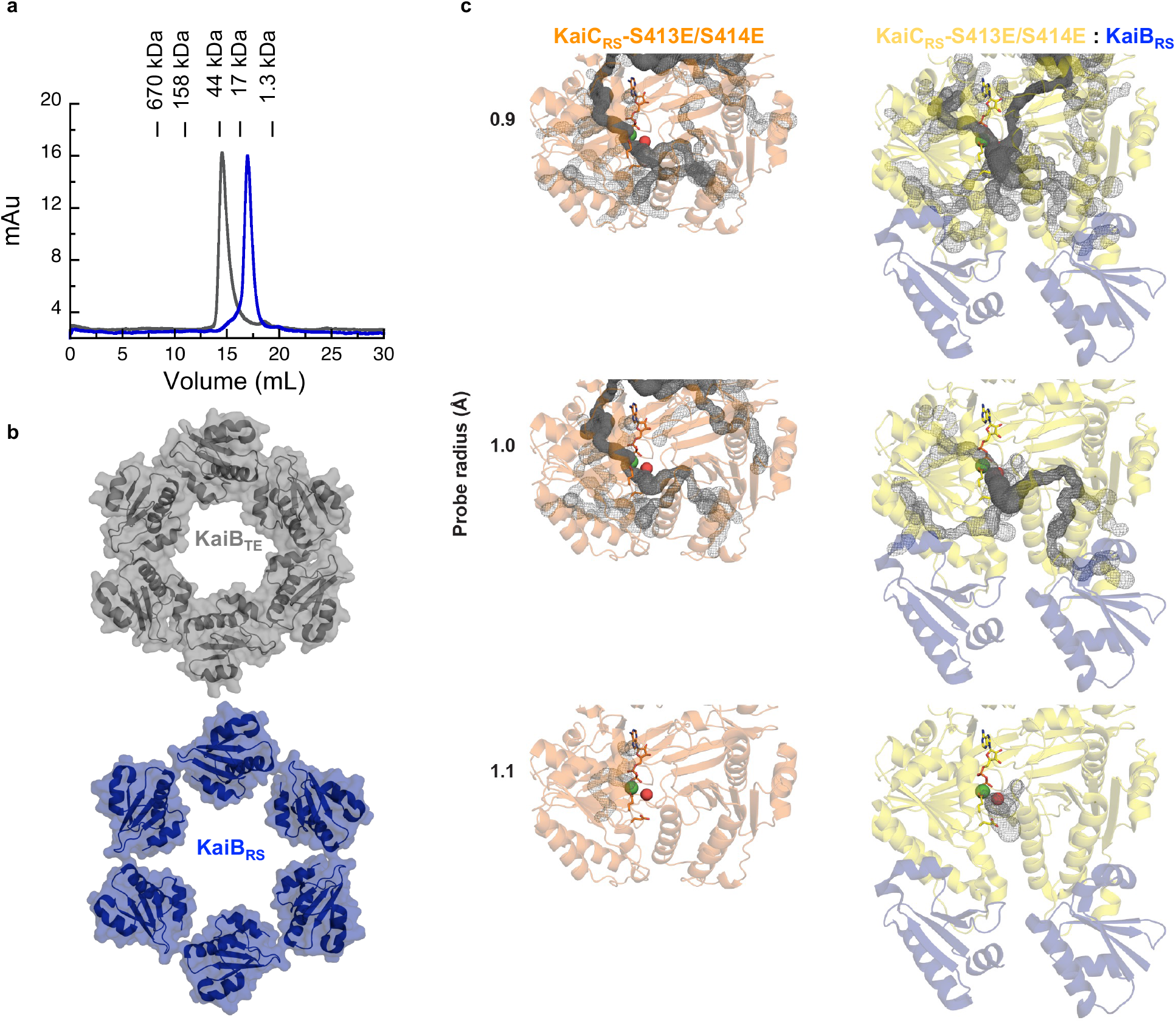
KaiB-KaiB interface in the KaiBC_RS_ complex affects the solvent accessibility into the active site of KaiC_RS_-CI. (**a**) Size-exclusion chromatography of KaiB_RS_ (blue) shows that it is monomeric in solution in contrast to KaiB_SE_ (gray), which elutes as a tetramer. Molecular-weight standards are shown above the chromatogram. **(b)** Structural comparison of KaiB_TE_ (gray, PDB 5jwq^26^) and KaiB_RS_ (blue) when bound to their corresponding KaiC hexamers. The PISA software package^37^ determines that for the KaiBC_TE_ complex the interface between the KaiB_TE_ monomers is 255 Å^2^, whereas the average interface between KaiB_RS_ monomersis only 45 Å^2^ in the KaiBC_RS_ complex. **(c)** To understand how KaiB_RS_ binding to KaiC_RS_-CI domain increases the hydrolysis rate, we investigated whether conformational changes modulated substrate access to the active site. The CAVER software^39^ was used to calculate tunnels (gray mesh) leading into the active site of the CI domain of KaiC_RS_-S413E/S414E alone (orange) and the KaiC_RS_-S413E/S414E:KaiB_RS_ complex (yellow:blue) with varying probe radii. In both structures, the active site was occupied by ADP:Mg^2+^ (sticks and green sphere, respectively). The crystal structure of bovine F1-ATPase in complex with a transition-state analogue (PDB 1w0j, chain D)^40^ was used as a reference to determine the position of the catalytic water molecule in the active site (shown as a red sphere). The calculated tunnels connect bulk solvent to the catalytic water when KaiB_RS_ is bound for probe radii larger than the default value of 0.9 Å, but never in its absence. These results suggest that KaiB_RS_ facilitates the access of water into the active site of KaiC_RS_-CI via long-range conformational changes and thus enhances ATP hydrolysis.

**Extended Data Fig. 10.**
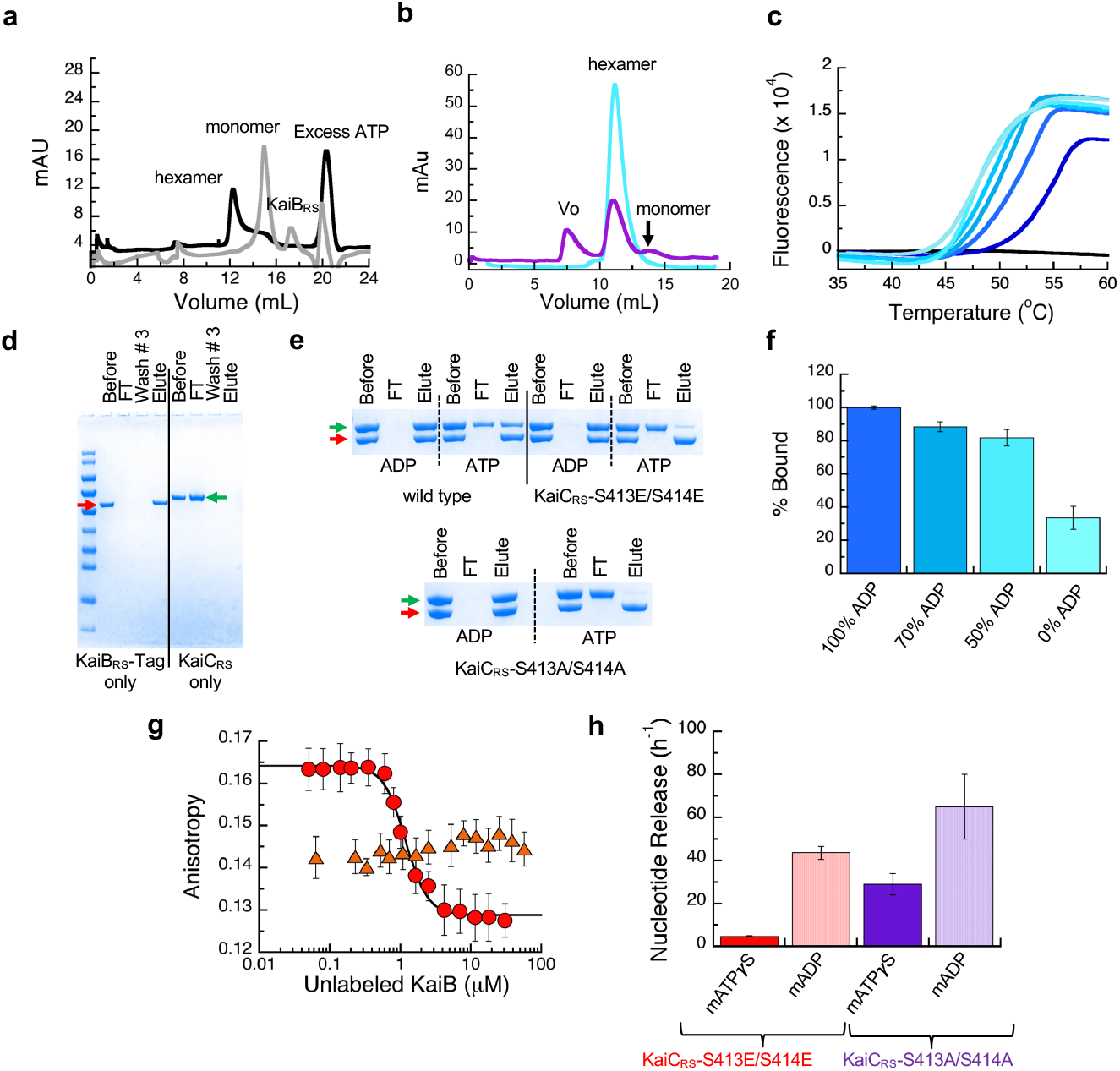
KaiB binds preferentially to the post-hydrolysis state of KaiC _RS_ and affects its stability. **(a)** Size-exclusion chromatography of 50 μM KaiC_RS_-CI (CI_RS_ domain) in the absence (black line, hexamer) and presence (gray line, monomer) of 50 μM KaiB_RS_ in 1 mM ATP buffer. **(b)** Size-exclusion chromatography of 50 μM KaiC_RS_-Δcoilin the presence of 50 μM KaiB_RS_ (purple). The reference sample (50 μM KaiC_RS_-Δcoil) is a hexamer in solution (cyan) and after the addition of 50 μM KaiB_RS_ the mixture was incubated at 30 °C for 3.5 h (purple) before running the samples again ona Superdex-200 10/300 GL column at 4 °C. These data show that binding of KaiB_RS_ results in (i) disassembly of the hexameric KaiC_RS_-Δcoil structure into its monomers and (ii) aggregation as detected by the elution in the void volume of the column (v_0_). **(c)** Thermal denaturation profiles for KaiC_RS_-S413E/S414E in the presence of 1 mM ADP are shown from dark to light blue for increasing concentrations of KaiB_RS_ (between 0 – 4 μM). The black line represents KaiB_RS_ alone (15 μM), which shows no fluorescence signal as it does not bind to SYPRO Orange due to a lack of a hydrophobic core. The T_m_ decreases upon the addition of KaiB_RS_, indicating that binding of KaiB_RS_ destabilizes the KaiC_RS_ dodecamer. Likely due to loosening up of interface and the KaiC_RS_ structure, thereby allowing for the formation of a tunnel that connects bulk solvent to the position of the hydrolytic water in the active site (see Extended Data Fig. 9). **(d)** SDS-PAGE analysis showing the control experiment for pull-down assay. The first four lanes after the molecular weight marker are KaiB_RS_-Tag samples (red arrow) and show that KaiB_RS_-Tag binds tightly to the column. The last four lanes are control pull-down assay experiments for KaiC_RS_ (green arrow) and show that KaiC_RS_ alone is unable to bind to the column. The lanes represent the initial sample used in pull-down assay (Before), flow-through after loading sample onto the column (FT), flow-through after washing the column three times with the binding buffer (Wash #3), and sample after elution with imidazole (Elute). (**e**) SDS-PAGE analysis of pull-down assay to measure the complex formation between KaiB_RS_-Tag and wild-type KaiC_RS_, KaiC_RS_-S413E/S414E, or KaiC_RS_-S413A/S414A in the presence of 4 mM ADP or ATP (with an ATP-recycling system). **(f)** Percentage of wild-type KaiC_RS_ bound to KaiB_RS_-Tag protein for different ATP-to-ADP ratios (4 mM total nucleotide concentration) at 25 °C as measured from pull-down assays. **(g)** Fluorescence anisotropy at 30 °C of unlabeled KaiB_RS_ competitively replacing the fluorophore-labeled KaiB_RS_ (KaiB_RS_-6IAF) from KaiC_RS_-S413E/S414E in the presence of 4 mM ADP (red circles, *K*_D_ value of 0.79 ± 0.06 μM) and 4 mM ATP with an ATP-recycling system (orange triangles). In the latter experiment no change in anisotropy is observed, indicating that only a small fraction of KaiB_RS_-6IAF is bound under these conditions. The average anisotropy and standard error were calculated from ten replicate measurements. **(h)** The mant-ATPγS or mant-ADP release is shown as bar graphs with observed rates of 4.8 ± 0.2 h^-1^ and 43.6 ± 3.0 h^-1^ for mant-ATPγS and mant-ADP releasing from KaiC_RS_-S413E/S414E, respectively, and 21.0 ± 3.0 h^-1^ and 65 ± 15 h^-1^ for mant-ATPγS and mant-ADP releasing from KaiC_RS_-S413A/S414A, respectively. The result shows that CII domain of KaiC_RS_ prefers binding of ATP over ADP. Experiments in panels f and h were performed in triplicate with error bars representing SD.

**Extended Data Table 1.** X-ray crystallography data collection and refinement statistics.

|  | KaiCRS<br>PDB: 8dba<br>(oligomeric state:<br>dodecamer) | KaiCRS-Δcoil<br>PDB: 8db3<br>(oligomeric state:<br>hexamer) |
| --- | --- | --- |
| **Data collection** |  |  |
| Space group | P1 | C222<sub>1</sub> |
| Cell dimensions |  |  |
| *a*, *b*, *c* (Å) | 105.11, 136.09, 146.2 | 94.06, 197.28, 150.19 |
| $\alpha$, $\beta$, $\gamma$ (°) | 93.13, 94.43, 108.09 | 90, 90, 90, |
| Resolution (Å) | 47.65 - 3.5 (3.625 - 3.5)* | 53.89 - 2.9 (3.004 - 2.9)* |
| *R*<sub>sym</sub> or *R*<sub>merge</sub> | 0.2213 (1.372) | 0.4267 (2.994) |
| *I* / σ*I* | 2.57 (0.73) | 5.54 (1.15) |
| Completeness (%) | 97.62 (94.08) | 99.89 (99.97) |
| Redundancy | 1.9 (2.0) | 7.2 (7.3) |
| **Refinement** |  |  |
| Resolution (Å) | 3.5 | 2.9 |
| No. reflections | 94151 (9047) | 31344 (3102) |
| *R*<sub>work</sub> / *R*<sub>free</sub> | 0.2323/ 0.2775 | 0.2308/0.2892 |
| No. atoms | 47510 | 10293 |
| Protein | 46772 | 10131 |
| Ligand/ion | 948 | 231 |
| Water | 67 | 0 |
| *B*-factors |  |  |
| Protein | 102.48 | 56.06 |
| Ligand/ion | 89.08 | 44.62 |
| Water | 72.52 | - |
| R.m.s. deviations |  |  |
| Bond lengths (Å) | 0.002 | 0.004 |
| Bond angles (°) | 0.62 | 0.73 |

\*Values in parentheses are for highest-resolution shell.
A single crystal was used for each structure.

**Extended Data Table 2.**
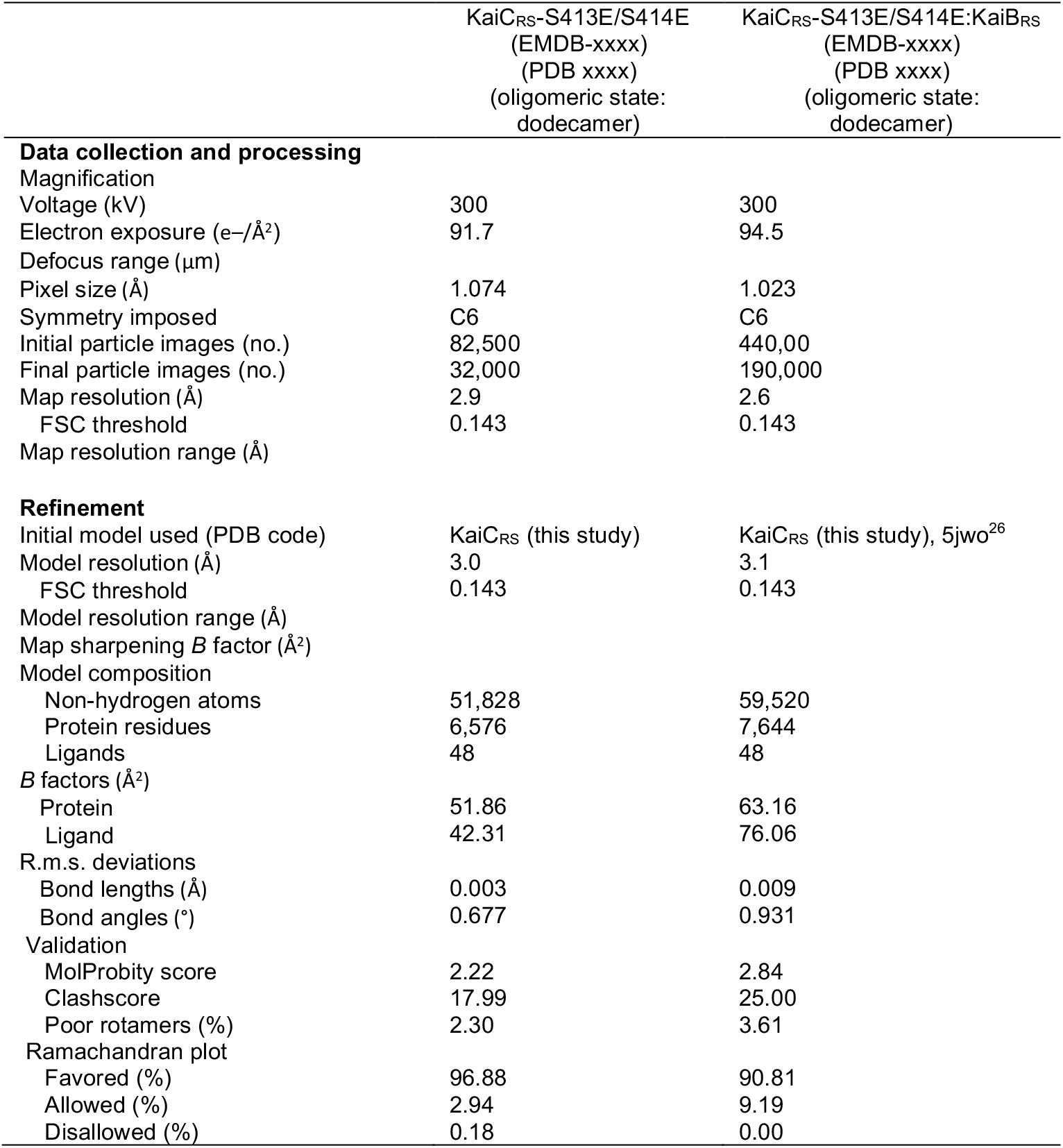
Cryo-EM data collection, refinement, and validation statistics.

**Extended Data Table 3.**
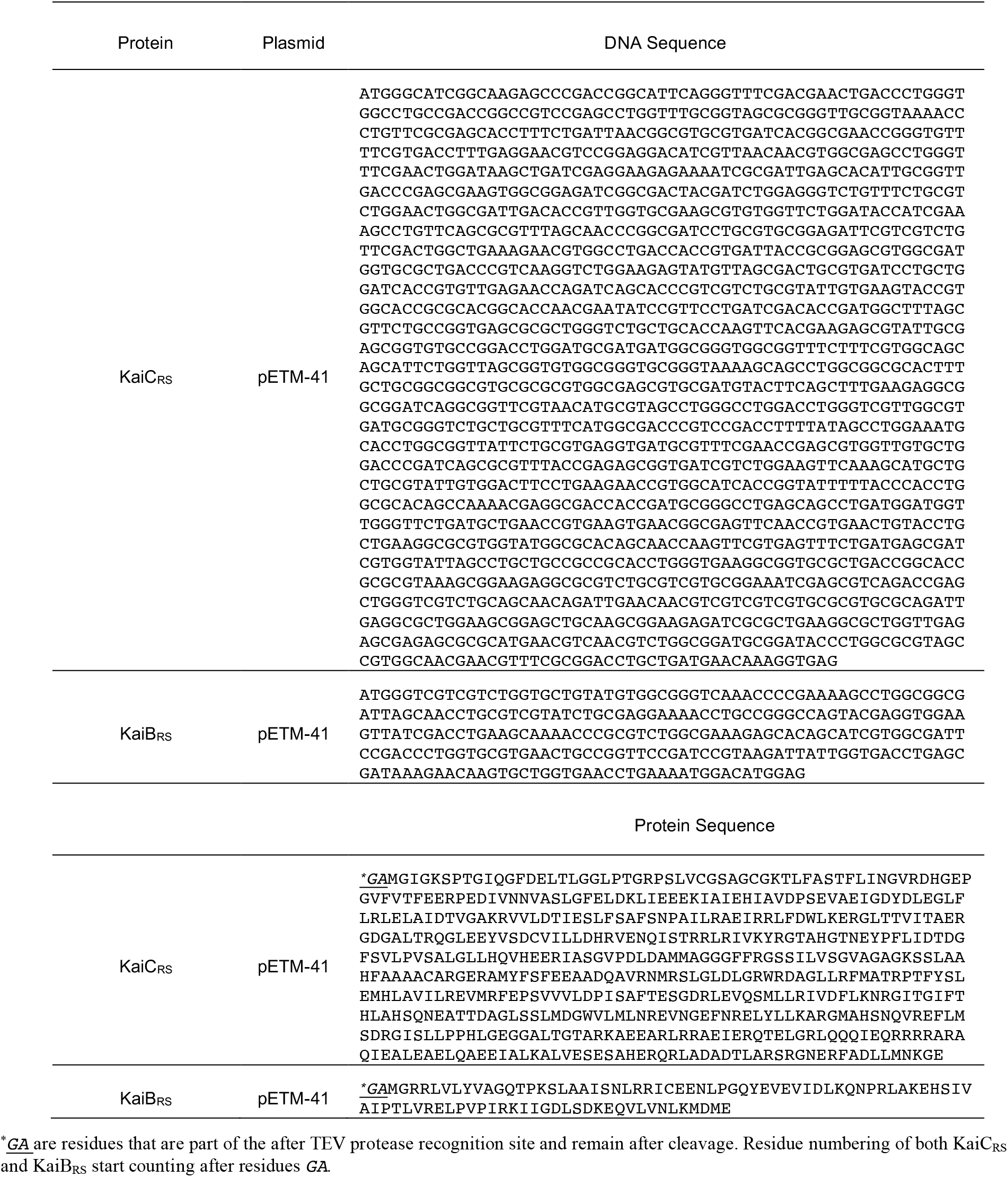
Codon-optimized DNA sequence for KaiC_RS_ and KaiB_RS_ constructs.

**Extended Data Table 4.**
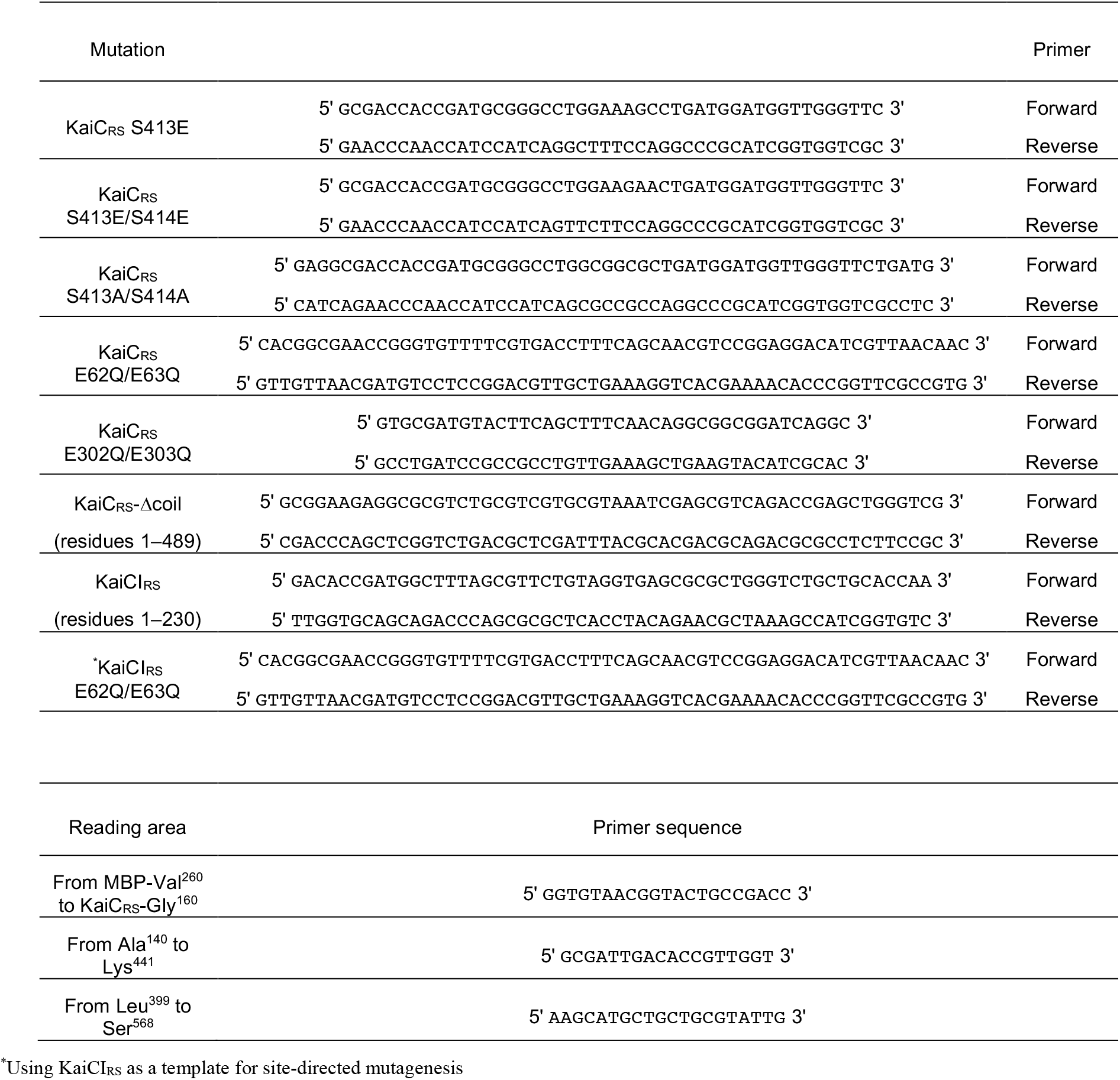
Primers for site-directed mutagenesis and sequencing of KaiC_RS_.

